# Comparative study on chromatic acclimation in the two *Synechococcus* species WH8109 and RS9916

**DOI:** 10.64898/2026.10.21.760676

**Authors:** Jeegna Solanki, Miguel Fernandez, Mika Pawitsch, Michelle M. Gehringer, Susanne Zehner, Nicole Frankenberg-Dinkel

## Abstract

Marine *Synechococcus* cyanobacteria can modify their light-harvesting phycobilisome antennae in response to spectral changes through chromatic acclimation (CA). CA type IV (CA4) involves adjustment of the ratio of the blue-absorbing phycourobilin (PUB) to the green-absorbing phycoerythrobilin (PEB). Although CA4 is associated with a genomic island occurring in two configurations (CA4-A and CA4-B), some strains carrying this island, including *Synechococcus* sp. WH8109, have been reported to lack CA4 activity. Here, we investigated whether WH8109 undergoes CA4 under blue light (BL) and green light (GL) conditions, using the CA4-positive strain RS9916 as a reference.
Whole-cell fluorescence, flow cytometry, isolated phycobilisome analysis, and sequencing of the CA4-B genomic island were used to assess pigment acclimation and identify potential genetic differences.
WH8109 showed a stable PUB:PEB ratio of approximately 0.71 under BL and GL, with no detectable changes in PUB content or phycobilisome chromophore composition. In contrast, RS9916 exhibited clear CA4, with PUB:PEB ratios of 1.57 under BL and 0.72 under GL. Sequencing revealed no insertions, deletions, or other sequence anomalies in the WH8109 CA4-B genomic island compared to the public sequence.
These results demonstrate that WH8109 does not undergo detectable CA4 under the conditions tested, despite possessing an intact CA4-B genomic island, highlighting a genotype–phenotype discrepancy that warrants further investigation.

**One sentence summary of the key message of the paper:** The original isolate of *Synechococcuss* sp. WH8109, obtained from the Roscoff culture collection, was confirmed to not undergo chromatic acclimation type-IV under changing blue-green light conditions.

**Graphical Abstract:** Upon acceptance of the manuscript, the authors should be prepared to submit a self- explanatory graphical abstract to the editorial office and to deposit the original data in a data repository or archive and provide the persistent identifier (eg DOI). This is an essential requirement before the manuscript can be forwarded to production.

## Introduction

Marine *Synechococcus* are abundant and ecologically important marine cyanobacteria whose success across diverse oceanic environments is partly associated with their ability to adapt their light-harvesting systems to prevailing light conditions (Chen et al., 2026; Flombaum et al., 2013; Mattei et al., 2025; Wang, Li, Jing, & Qin, 2022). Their primary light-harvesting antennae, the phycobilisomes (PBS), contain phycobiliproteins (PBP) carrying bilin chromophores that absorb light over different regions of the visible spectrum (Kehoe et al., 2025; Sanfilippo, Garczarek, Partensky, & Kehoe, 2019; Wang et al., 2022), accounting for their wide distribution from ocean surface waters to depths of 150-200 m (Grébert et al., 2022), with occasional reports of *Synechococcus* occurrence of between 200 – 600 m (Lindell & Post, 1995). Competition for light has resulted in the evolution of three major pigment types (PT) within the marine *Synechococcus* cyanobacteria (Humily et al., 2014; Six et al., 2007). Blue light (BL) specialists, PT 3c, favour a relatively high amount of the BL absorbing phycourobilin (PUB) to the green light absorbing phycoerythrobilin (PEB) on the phycobilisome rods. Green light (GL) specialists, PT 3a, on the other hand, have a lower PUB:PEB ratio (Dufour et al., 2025; Kehoe, 2010; Kehoe et al., 2025; Sanfilippo, Garczarek, et al., 2019). The third pigment type, PT 3d, encompasses strains capable of conducting chromatic acclimation type IV (CA4), a marine *Synechococcus* acclimation strategy in which the PUB ratio of the PBS is adjusted in response to ambient BL- and GL-conditions to optimise photon collection. Under BL, increased incorporation of PUB (*λ*max ≈ 495 nm), facilitated by the action of a lyase isomerase enzyme, enhances absorption in the blue region, whereas GL favours increased incorporation of PEB (*λ*max ≈ 550 nm) (Dufour et al., 2025; Grébert et al., 2021; Humily et al., 2014; Sanfilippo, Nguyen, et al., 2019; Shukla et al., 2012). Unlike other forms of chromatic acclimation, CA4 primarily involves changes in bilin chromophore composition rather than major changes in the abundance of the PBP themselves. Thus, changes in the PUB:PEB ratio, Exc495:545, recorded as the fluorescence of PEB at 570 nm, from excitation at 495 nm (PUB) and 545 nm (PEB), provide a measurable indicator of CA4 activity (Grébert et al., 2021; Humily et al., 2014).

CA4 strains have two genetically distinct pigment types, PT 3dA and PT 3dB, associated with the genomic islands CA4-A and CA4-B, respectively (Humily et al., 2014; Six et al., 2007). Both configurations encode components required for the light-dependent modification of PBS chromophores, although the islands differ both genetically and structurally (Figure 1) (Grébert et al., 2021; Sanfilippo, Nguyen, et al., 2019; Shukla et al., 2012). Recently it was shown that PT 3dA and PT 3dB strains required different blue to green light ratios to trigger the process of CA4 (Dufour et al., 2024).

**Figure 1.**
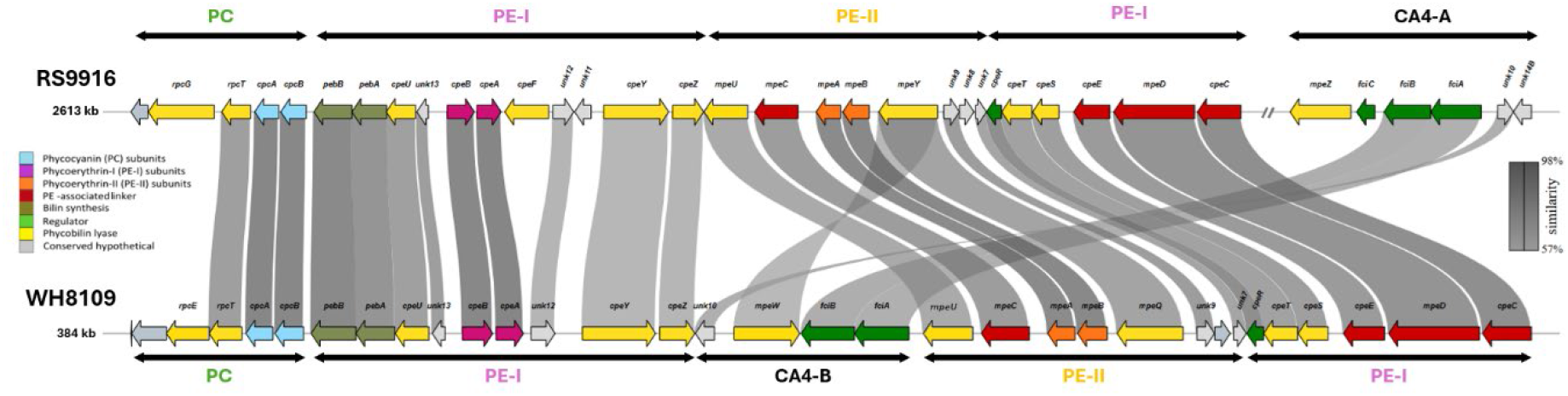
Organisation of the phycobilisome (PBS) rod genomic region of *Synechococcus* sp. RS9916 (top) and WH8109 (bottom) and their CA4 islands. The schematic representation of the PBS region was generated using sequence data from <u>NCBI</u> (GCF_000153825.1 and NZ_CP006882.1, for RS9916 and WH8109, respectively) and <u>Cyanorak</u> (Garczarek et al., 2021). The CA4-A (RS9916) and CA4-B (WH8109) regions are marked with black text, genes required for production of PC labelled in green text, phycoerythrin-I (PE-I) in pink text and phycoerythrin -II (PE-II) in yellow text. Yellow coloured open reading frames (ORFs) are phycobiliprotein lyase genes, teal coloured genes represent PC α- and β-subunits, pink coloured genes indicate PE-I α-and β-subunits, orange coloured genes are PE-II α- and β-subunits, red coloured genes encode linker proteins, while genes with unknown function (*unk*) are coloured grey. Green coloured open reading frames encode regulatory proteins (Kehoe et al., 2025). Genes are coloured according to their inferred function. Comparison was made via BlastP with the translated RS9916 genomic region as query identity and a minimum of 40% similarity, similarity levels are showed in shades of grey ranging from 57% to 98%. Both image and Blast were produced with GeneViewer R (van der Velden, 2025). Note: While Cyanorak has *mpeQ* annotated as *mpeY*, we followed the suggestion of Gebert et al (Grébert et al., 2021) renaming *mpeY* from PT3dB strains to *mpeQ.* Details see also under Material and Methods.

Grebert et al., suggested that CA4-A strains naturally exhibit a GL specialized phenotype (defined by a low PUB:PEB ratio) (Grébert et al., 2021). CA4 activation allows these strains to modify their spectral characteristics under BL by inducing the expression of a phycobilin lyase-isomerase gene, *mpeZ*, to increase the amount of PUB incorporated in the PBS. CA4-B strains, on the other hand, have a baseline specialisation for BL (high PUB:PEB ratio), and CA4 gives them the ability to adapt to GL by inducing expression of the PEB lyase gene, *mpeW*, thereby increasing the amount of PEB in the light harvesting PBS.

The presence of a CA4 island, however, does not necessarily predict the observed acclimation phenotype. This is particularly relevant for *Synechococcus* sp. WH8109, which carries a CA4-B genomic island (Figure 1) but has yielded conflicting reports regarding its ability to undergo CA4 (Humily et al., 2014; Lovindeer, Abbott, Medina, & Mackey, 2021). Humily et al. reported that WH8109 maintained a relatively constant PUB:PEB ratio under BL and GL and therefore did not exhibit CA4 (Humily et al., 2014).

In contrast, Lovindeer et al. subsequently reported a light-dependent increase in the PUB ratio under BL and a decrease under GL, consistent with CA4 activity (Lovindeer et al., 2021). Resolving this discrepancy is important for understanding the relationship between the presence of the CA4-B genomic island and its functional expression in WH8109.

Here, we investigated the capacity of *Synechococcus* sp. WH8109 to undergo CA4 under BL and GL conditions using *Synechococcus* sp. RS9916, a PT 3dA, CA4-A capable, strain, as a reference. Changes in cellular PUB:PEB ratios were assessed in cultures, and individual cells, and the chromophore composition of isolated PBSs was examined to determine whether changes detected at the whole-cell level reflected changes in the bilin composition of the light-harvesting antenna. In addition, the CA4- B genomic island of WH8109 was sequenced to assess whether sequence alterations could account for differences between the previously reported CA4 phenotypes.

## Materials and Methods

### Materials and Reagents used

All chemicals used were reagent grade or better and purchased from Merck, AppliChem and Roth unless otherwise mentioned. Centrifugation steps were performed with an Eppendorf AG 5415D unless otherwise mentioned.

### Strains and growth conditions

Non-axenic culture of *Synechococcus* sp. WH8109, originally isolated from the Sargasso Sea, northwest Atlantic ocean in 1981 by L. Brand (Waterbury, 1986) was kindly provided by Prof. Debbie Lindell (Technion-Israel Institute of Technology, Haifa, Israel) and is referred to as ‘WH8109’ in this study. Additionally, the CA4-A strain *Synechococcus* sp. RS9916, isolated from the Gulf of Aqaba in the Red Sea in 1999 (Fuller et al., 2003) and the original *Synechococcus* sp. WH8109, henceforth referred to as ‘WH8109_RC’, were purchased from the Roscoff culture collection (https://roscoff-culture-collection.org/rcc-strain-details/2033). Strains were grown at 21 ± 2°C at 20 ± 2 µmol photons m^-2^s^-1^ continuous light (measured using a Li-Cor Quantum sensor and Li-250A Light Meter, Li-Cor, USA), in artificial sea water (ASW) + NO3 medium (Roscoff Culture, 2019)(Roscoff Culture, 2019) in 250 ml of acid washed flask (three baffled flask, Duran) with 120 rpm shaking. In some experiments, WH8109 cultures were grown in sea water nutrient (SN) medium as described (Lovindeer et al., 2021). Growth was monitored through chlorophyll a (Chl-*a*) fluorescence measurements and used to determine the growth rates under the different culture conditions tested.

### Light acclimation experiment

Experimental cultures were grown in a custom-made culture apparatus whereby each culture bottle was individually illuminated by an adjustable light source (Supplementary Figure 1 A) set at 20 ± 2 µmol photons m^-2^s^-1^, 21 ± 2°C for each light investigated using LEE lighting filters (BL: dark blue, #119, 455 nm peak transmission; GL: primary green, #139, 525 nm peak transmission; WL: full spectrum white light LED). Light spectra and intensities were confirmed (Supplementary Figure 1 B) using an optical multichannel analyser (USB2000^+^ Ocean Optics, Germany). Strains were inoculated in triplicate from exponentially growing cultures from white light conditions, at a Chl-*a* fluorescent intensity of 100 units at 680 nm (440 nm excitation) in ASW media and exposed to continuous BL or GL at 20 ± 2 µmol photons m^-2^s^-1^, 21 ± 2°C, with shaking at 120 rpm (ELMI DOS-20L, Latvia). Cultures were diluted to 100 Chl-*a* fluorescent units mL^-1^ after eight days (WH8109) or ten days (RSS9916) and transferred to the alternative light source for an additional acclimation period. Growth and pigment content were assessed daily through fluorescence measurements. Chromatic acclimation was determined to be complete when no further changes were observed in PUB:PEB fluorescence ratio.

Lovindeer et al. (Lovindeer et al., 2021) conducted CA4 light acclimation experiments on *Synechococcus* WH8109 growing in SN medium, that demonstrated CA4 activity. To eliminate potential effects of medium on CA4 induction, we repeated the light exposure experiments on WH8109 grown in SN medium (Lovindeer et al., 2021). To exclude the possibility of our WH8109 strain having undergone a genetic modification to eliminate the proposed CA4 activity previously observed, we obtained the strain directly from the same source, the Roscoff Culture Collection (Lovindeer et al., 2021). The above acclimation experiment was repeated using the newly purchased stock of WH8109_RC. We additionally sequenced the CA4 island of our original WH8109 strain to confirm the lack of any mutations (See section 2.7).

### Fluorescence and absorbance measurements

*In vivo* Chl-*a* fluorescence was measured at room temperature by exciting whole cell culture material at 440 nm and recording the emission from 650 to 750 nm (Fluorescence spectrometer FP-8300, Jasco, Germany). Fluorescent readings at 680 nm (Supplementary Figure 2A) were used to calculate the growth rates (Supplementary Figure 2B).

The *in vivo* PUB and PEB fluorescence of culture material collected in parallel was recorded by measuring the emission fluorescence at 570 nm, off an excitation scan from 440 to 560 nm (FP-8300 Jasco, Germany) and plotted after normalizing against Chl-a fluorescence (Supplementary Figures 3, 4 & 5). The PUB:PEB (Exc495:545) ratio, was calculated from the 570 nm emission values, arising from excitation at 495 nm (PUB) and 545 nm (PEB) (Dufour et al., 2024; Grébert et al., 2021; Lovindeer et al., 2021). Selected bands obtained from the PBS density gradient isolations (Section 2.6) were analysed using UV-Vis spectroscopy (8453 UV-Visible system; Agilent). The absorption of each fraction indicated in Figure 4, as well as Supplementary Figures 7, 8, 9 and 10 were recorded between 300 nm and 800 nm (8453 UV-Visible System of Agilent) in the dark at RT, using 0.75 M potassium phosphate buffer as reference.

### Flow cytometric analysis

Flow cytometric analysis (FCM) is commonly used to measure chromatic acclimation in *Synechococcus* species by tracking changes in the fluorescence signatures of BL and GL absorbing phycobiliproteins, specifically the PUB:PEB ratio, as well as cellular phycoerythrin (PE) fluorescence (Chisholm et al., 1988; Corzo, Jimenez-Gomez, Gordillo, & Niell, 1999; Dufour et al., 2024; Dufour et al., 2025; Tanaka, Hashihama, & Katano, 2024). *Synechococcus spp.* RS9916 and WH8109 cultures were analysed on day 0, 4 d, 8 d and 10d, to monitor chromatic acclimation. Samples were diluted to a concentration below 1 x 10^6^ cells per mL prior to analysis. The flow rate was set to 25 μL min^-1^ and a minimum of 6.8 x 10^5^ cells were analysed on an Attune NxT flow cytometer (Fisher Scientific). Fluorescence was detected using laser excitation at 488 nm and detection in channel BL2 (574 nm) and BL3 (695 nm) corresponding to PE and Chl-*a*, respectively.

### Phycobilisome isolation

Phycobilisomes were isolated from culture material of WH8109 and RS9916 after exposure to either BL or GL (Heck et al., 2024), to confirm the PUB:PEB (Exc495:545) results. Biomass was harvested from 150 ml late exponential phase (day 15) cultures by centrifugation at 10,000 x *g* for 20 min at RT, at a Chl-a fluorescence of ∼300 units. Pellets were resuspended in 0.25 ml of 0.75 M potassium phosphate buffer (pH 7), and centrifuged at 4000 x *g* for 5 min. The pellet was mixed with 0.1 mm FastPrep silica beads (BioSpec, USA) in 1:1 ratio, resuspended in 0.5 ml of 0.75 M potassium phosphate buffer (pH 7) and the cells disrupted in a bead beater (MM400 mixer mill, Retsch, Germany) for 5 cycles of 1 min at 30 beats per second, at 1 min intervals. The samples were subsequently centrifuged twice at 15 000 x *g* for 15 min at RT, to remove cellular debris and silica beads. TritonX-100 was added to the cleared lysates to a final concentration of 2% (v/v) and the samples again centrifuged at 15000 x *g* for 15 min at RT. The protein content of the ensuing phycobilisome containing lower phase was determined using the Bradford Assay (Roti-Quant, Carl Roth, Germany) (Bradford, 1976). Concentrations were calculated from a BSA standard curve (R^2^ = 0.99). Around 1.5 - 2 mg protein of the PBS containing fractions were loaded onto a discontinuous sucrose gradient consisting of 1.5 M, 1 M, 0.75 M and 0.5 M sucrose in 0.75 M potassium phosphate buffer (pH 7). The loaded gradients were then centrifuged 13000 x *g* for 16 h at 20°C (Optima MAX-XP, Beckman Coulter) in a MLS-50 swinging-bucket rotor.

Fractions obtained from the sucrose gradient were characterised by UV–Vis spectroscopy and fluorescence spectral analysis, as described above (Section 2.4). Additionally, proteins were precipitated from the sucrose gradient fractions deemed to contain complete PBSs. Acetone was added to a final concentration of 80% acetone, mixed thoroughly and incubated overnight at -20°C. The samples were subsequently centrifuged at 13 000 x *g* at 4°C to pellet the purified protein while leaving the remaining sucrose in the supernatant. The proteins were resuspended in 0.5 M potassium phosphate buffer (pH 7). Between 5–15 μL of the purified protein samples were mixed with 4 x SDS sample loading dye and heated at 95°C for 5 min, before being loaded onto a gradient SDS-PAGE of 4% and 20%. Protein Standard (NEB P7717) was run in parallel to determine the size of the proteins, after visualisation using Coomassie Blue G-250 staining (Supplementary Figure 11).

### Comparative genomic analysis of the phycobilisome region

Genomic sequences of the CA4-B island from *Synechococcus* sp. WH8109 (NZ_CP006882.1) and *Synechococcus* sp. RS9916 (NZ_DS022299.1) were retrieved from NCBI. Gene annotations were obtained from CyanoRAK(Garczarek et al., 2021). The gene *mpeQ*, annotated as *mpeY* in CyanoRAK, was designated *mpeQ* according to the functional characterization by (Grébert et al., 2021). Sequence similarity between the two genomic regions was assessed using BLASTP and the derived amino acid sequences, with the RS9916 region as the query and a minimum sequence identity threshold of 40%. The resulting similarity analysis and genomic region visualization were generated using the GeneViewer R package (van der Velden, 2025). Sequence similarities are represent ed by grey shading in Figure 1. Additional labels identifying specific regions were added using Microsoft PowerPoint.

### Protein structure prediction and comparison

The structures of the hypothetical proteins MpeW, MpeZ, and MpeY were predicted using the ESMFold protocol via the Colab research platform (Mirdita et al., 2022). The predicted structures were subsequently compared and visualized using PyMOL version 3.1.9 (Schrödinger, 2021).

### Statistical analysis

Experiments were conducted in triplicate and the data is potted as the mean values and standard deviation (± SD). PBS isolations and FCM were only conducted in duplicate. To ensure a normal distribution, the Kolmogorov-Smirnov test was employed (OriginLab). Subsequently, an F-test was executed for the analysis of variances, and the significance of the results was assessed through a two-tailed Student’s t-test (Microsoft Excel 2016). Analysis of Variance (ANOVA) was used to determine whether there was a significant difference in growth rates of *Synechococcus* WH8109 and *Synechococcus* RS9916 under each light condition. In addition, the F-test was used to determine variance of analysis and a two-sample T-test was used to calculate *p* values. Because there were four light conditions and more than one dataset, the *p*-value was corrected using the Bonferroni correction to reduce errors. Microsoft Excel 2016 was used for all calculations.

## Results

### Comparative analysis of the phycobilisome rod genomic region and CA4 genomic islands of *Synechococcus* sp. RS9916 and WH8109

Due to the conficting reports on the ability of *Synechococcus* sp. WH8109 to perform CA4, we reanalysed the PBS rod genoic region and associated CA4 genomic island and the physiolgical response of both strains to BL and GL. Both *Synechococcus* sp. RS9916 and WH8109 possess a CA4 genomic island that enables adjustment of the PBS PEB:PUB ratio in response to light colour. However, they carry distinct island configurations: RS9916 represents pigment type 3dA (CA4-A), with *fciA*, *fciB*, *fciC* and *mpeZ*, whereas WH8109 represents pigment type 3dB (CA4-B), with *fciA*, *fciB* and *mpeW*, but lacking *fciC* (Figure1). The two strains also differ in the associated chromophore-modification machinery: in RS9916, MpeZ promotes PUB attachment to Cys83 of the α-subunit of phycoerythrin II (MpeA) under BL, competing with the PEB lyase MpeY; in CA4-B strains, the corresponding functions are proposed to be performed by MpeQ (PUB attachment under BL) and MpeW (PEB attachment under GL), respectively. Thus, although both systems regulate chromophore composition at the same target site, they employ distinct enzyme pairs and differ in their regulatory organisation.

### Influence of BL and GL on the growth of S*ynechococcus* spp. WH8109 and RS9916

Chl-*a* is the primary pigment in photosynthesis, and measuring its *in vivo* fluorescence is a non-invasive, real-time method for tracking *Synechococcus* growth (Dufour et al., 2024; Lovindeer et al., 2021). The phototrophic growth of *Synechococcus* spp. WH8109 and RS9916, cultivated under BL, GL and WL, was tracked using Chl-*a* fluorescence emission measurements (Supplementary Fig. 2 A & B).

Both WH8109 and RS9916 exhibited the highest growth rates under WL, with 0.13 ± 0.06 Chl-*a* fluorescence units (FU). day^-1^ and 0.11 ± 0.06 Chl-*a* FU. day^-1^, respectively.

These growth rates matched those obtained under GL conditions (Supplementary Fig. 2 C). In comparison, BL cultures exhibited significantly reduced growth rates (p< 0.05) of 0.1 ± 0.05 Chl-*a* FU. day^-1^ for WH8109, and a non-significant, yet reduced growth rate of 0.09 ± 0.05 Chl-*a* FU. day^-1^ for RS9916 (Supplementary Fig.2 C).

**Figure 2.**
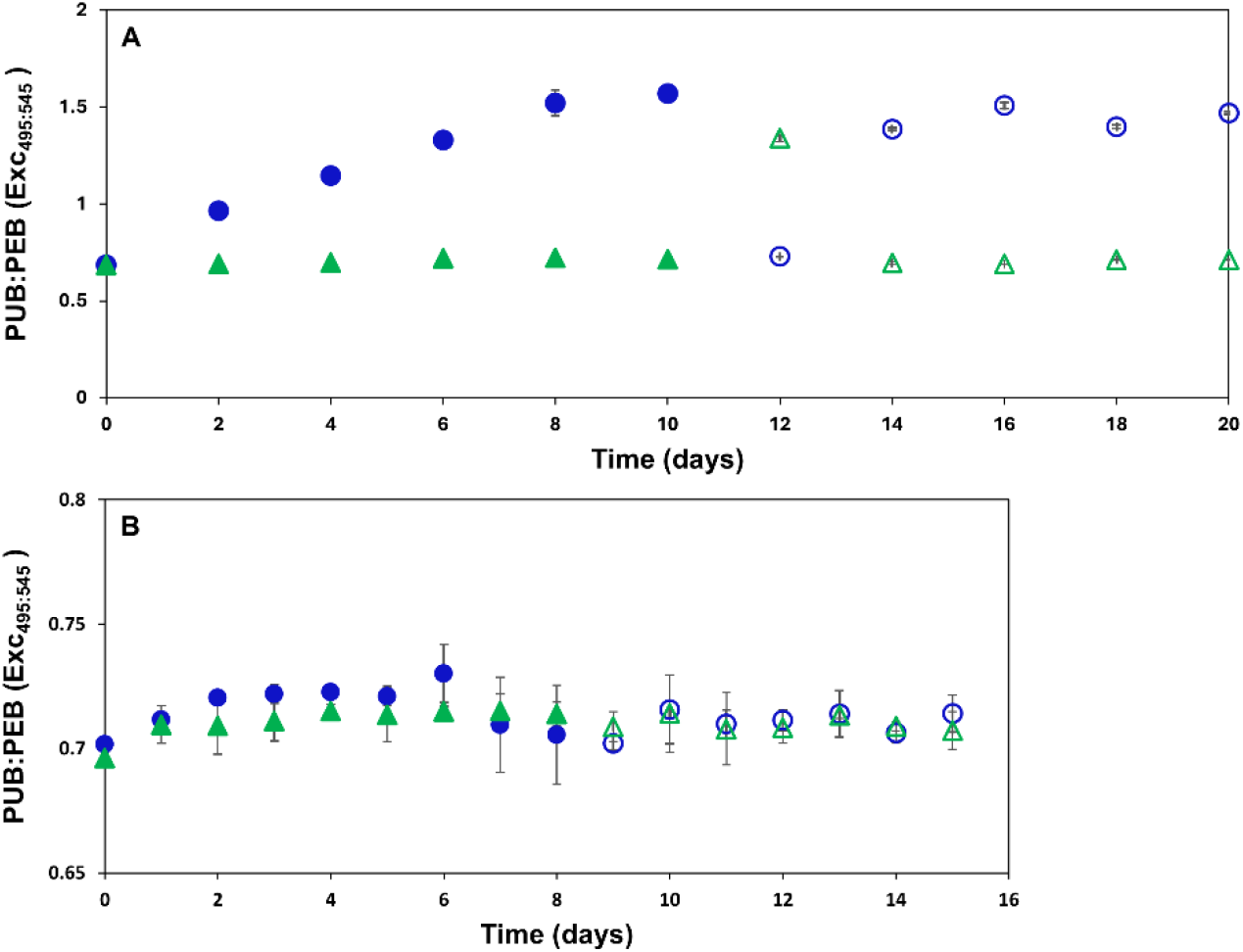
Comparative PUB:PEB (Exc_495:545_) *in vivo* fluorescence of *Synechococcus* spp. RS9916 and WH8109in ASW medium. Depicted are the average daily calculated PUB:PEB (Exc_495:545_) values during the light acclimation experiments. Ratios obtained from the primary BL or GL growth phase for **(A)** RS9916 (days 1-10) and **(B)** WH8109 (days 1-8) are plotted as blue dots or green triangles respectively. Open circles and triangles represent the average PUB:PEB ratios in the cultures after the light was changed to the alternative light colour. Error bars represent 1 standard deviation (SD) of triplicate cultures and are often too small to be visible.

### WH8109 does not exhibit a change in the PUB:PEB (Exc495:545) ratio as observed for RS9916 undergoing CA4

To confirm whether WH8109 could conduct CA4, we performed light acclimation experiments under BL or GL conditions and compared its PUB:PEB (Exc495:545) ratios to those from RS9916, a strain known to undergo CA4. The 570 emission spectra of whole cells indicated a clear shift in PUB content in RS9916 under BL conditions (Figure 2A, Supplementary Figure 3). The average PUB:PEB (Exc495:545) of 1.57 ± 0.02 was obtained on day 10 under BL, with it recorded as 1.45 ± 0.00 on day 20, after shifting the culture from GL to BL growth conditions (Table 1). These values were significantly raised compared to the PUB:PEB (Exc495:545) obtained under GL conditions. In contrast, no shift in PUB content was observed in the 570 nm spectra of WH8109 grown in ASW (Figure 2B, Supplementary Figure 3), neither were significant differences recorded for the PUB:PEB (Exc495:545) ratios (Table 1).

**Table 1.** PUB:PEB (Exc_495:545_) of *Synechococcus* spp. RS9916 and WH8109 under various light and medium conditions. Table listing the average PUB:PEB (Exc_495:545_) of cultures used in the different light acclimation experiments conducted to assess the potential of *Synechococcus* sp. WH8109 to conduct CA. The values for RS9916 were obtained from cultures grown in ASW medium under BL or GL for 10 days (Figure 2B). The average PUB:PEB (Exc_495:545_) values for WH8109 grown in ASW (Figure 2A) or SN medium (Supplementary Figure 6A), and for WH8109_RC grown in ASW (Supplementary Figure 6B) were obtained from measurements made on day eight for BL and GL grown cultures. The data shows the average and standard deviation of three independent biological replicates. *** indicates a significant difference of BL to GL grown cultures on the same day of p< 0.001.

| Strain | RS9916 | WH8109 | WH8109 | WH8109_RCC |
| --- | --- | --- | --- | --- |
| Medium | ASW | ASW | SN | ASW |
| BL | $1.57 \pm 0.02$ *** | $0.71 \pm 0.02$ | $0.71 \pm 0.02$ | $0.72 \pm 0.00$ |
| GL | $0.72 \pm 0.01$ | $0.71 \pm 0.01$ | $0.71 \pm 0.00$ | $0.71 \pm 0.01$ |

**Table 1. PUB:PEB (Exc<sub>495:545</sub>) of *Synechococcus* spp.**
| Strain | RS9916 | WH8109 | WH8109 | WH8109_RCC |
| --- | --- | --- | --- | --- |
| Medium | ASW | ASW | SN | ASW |
| BL | 1.57 ± 0.02 *** | 0.71 ± 0.02 | 0.71 ± 0.02 | 0.72 ± 0.00 |
| GL | 0.72 ± 0.01 | 0.71 ± 0.01 | 0.71 ± 0.00 | 0.71 ± 0.01 |

Culturing WH8109 in SN medium as reported by (Lovindeer et al., 2021), did not induce changes in the recorded 570 nm emission spectra (Supplementary Fig. 4), nor affect the PUB:PEB (Exc495:545) ratio when compared to WH8109 grown in ASW medium (Table 1, Supplementary Figure 6A).

In order to rule out any mutation in our WH8109 sp., we acquired a fresh culture from the Roscoff Culture Collection (WH8109_RCC). However, the newly acquired WH8109_RC did not, however, demonstrate any differences in its 570 emission spectra (Supplementary Fig. 5), nor its PUB:PEB (Exc495:545) ratio, when compared to WH8109 (Table 1, Supplementary Fig. 6B).

### Cellular PE fluorescence of WH8109 did not increase under BL

Fluorescence emission of PE was monitored by FCM for GL and BL grown *Synechococcus* sp. RSS9916 and WH8109 cells over an acclimation period of 10 d (days 0, 4, 8 and 10). Consistent with our previous findings, the strains RS9916 and WH8109 reacted differently to the GL and BL conditions. For confirmation, fluorescence PUB:PEB ratios were determined instantly before FCM analysis. During the acclimation period a clear increase of PE fluorescence per cell was observed for RS9916 in GL, and even more pronounced, in BL conditions (Figure 3). In contrast, in WH8109 the cellular PE fluorescence remained stable under both BL and GL incubation. All cultures displayed a uniform distribution of cell population. No hints for population heterogeneity were observed. Overall, the observations on cellular level obtained by FCM are consistent with the data of fluorescence spectroscopy on culture level.

**Figure 3.**
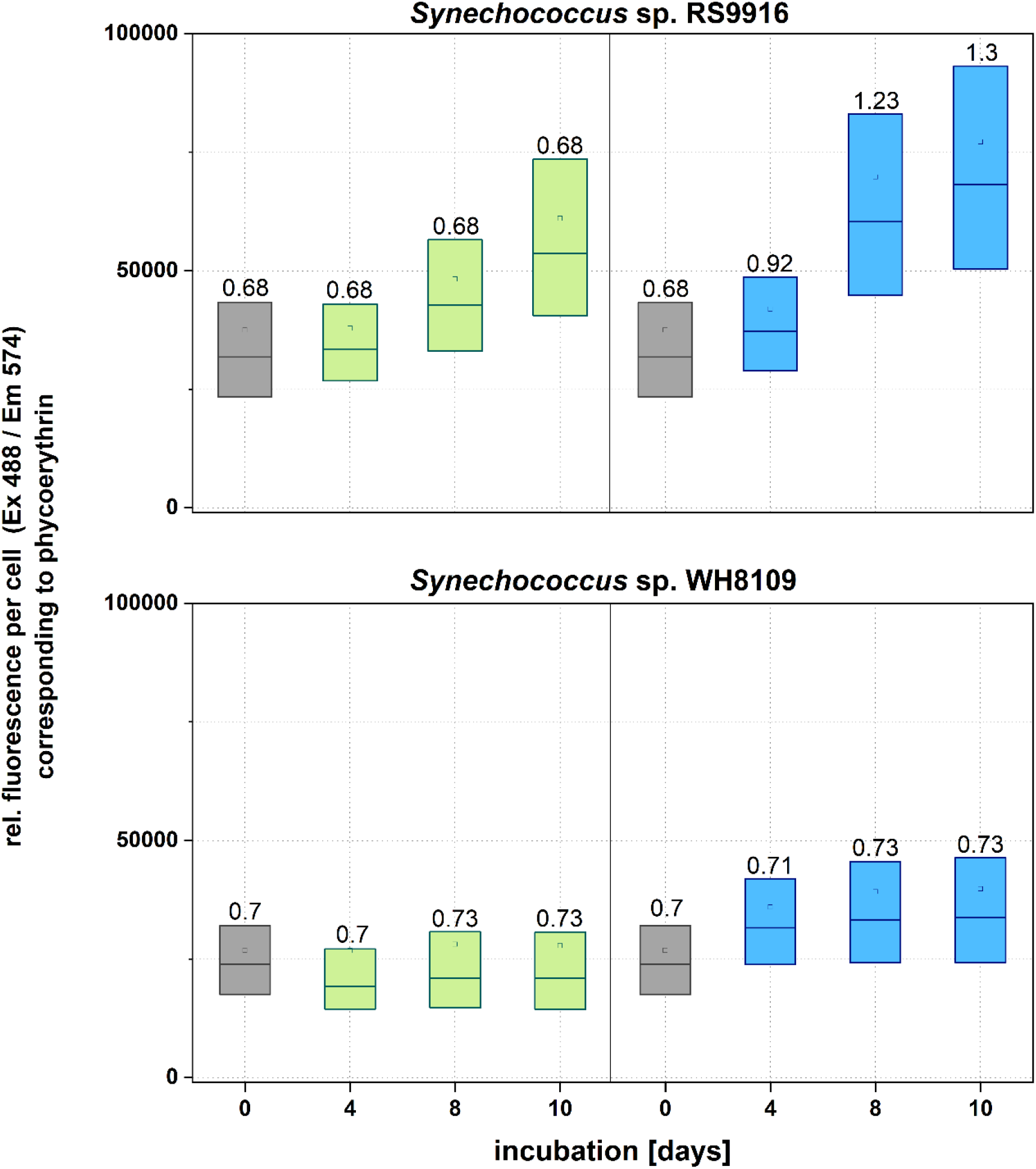
Flow cytometric PE fluorescence per cell of *Synechococcus* spp. RS9916 and WH8109 during acclimation to green and blue light over 10 days. Each box represents the mean of the fluorescence measurements per cell and the range of all cellular fluorescence values (25-75 %) performed for two individual cultures per strain in each light color condition (*n* = 2). The numbers above each box are PUB:PEB fluorescence ratio measured in the culture instantly before FCM analysis. Grey boxes indicate the fluorescence of white light grown cultures at the start of the acclimation experiments. Green and blue boxes depict the values for GL and BL irradiated cultures at the indicated day of the acclimation experiment, respectively. Light intensity during acclimation: blue light 9.5 μmol photons m^−2^ s^−1^; green light 8.5 μmol photons m^−2^ s^−1^ with 14/10 hours light-dark-cycle. Data analysis was performed with Origin 2022.

### Isolated phycobilisomes of WH8109 grown in BL do not show increased PUB:PEB (Exc495:545) ratios as seen for RS9916

To confirm that intact PBS reflected the whole cell fluorescence PUB:PEB ratios, PBS were isolated from cultures of RS9916 and WH8109 grown in ASW media with continuous illumination under BL, GL and WL conditions. The profiles of the PBS fractions obtained during sucrose gradient separation are presented for the positive CA4 acclimating strain RS9916 grown in BL (Figure 4A), GL (Supplementary Figure 7A) and WL (Supplementary Figure 8A). The sucrose gradients all exhibited similar separation patterns (Fig.4A, Supplementary Figures 7A and 8A). The yellowish pinkish fraction at 0.5 M, labelled as Fraction 1, was identified as Chl-*a*, due to its strong fluorescence at 680 nm after excitation at 440 nm (Supplementary Figures 7B and 8B). Fraction 2, located at 0.75 M sucrose, exhibited a UV-VIS spectrum with absorption peaks at the PUB λmax of 495 and the λmax of 545 nm for PEB (Figure 4C, Supplementary Figures 7C and 8C). Excitation of Fraction 2 at 495 nm yielded the distinctive fluorescent emission peaks for PE and PC (Figure 4E, Supplementary Figures 7D and 8D), indicating that this fraction contained complete PBS. The fluorescence scan of Fraction 2, recorded at 570 nm, provided the relative fluorescence peaks of PUB:PEB (Fig. 4G, Supplementary Figures 7E and 8E). The relative PUB:PEB ratio for the whole PBS is higher for BL grown cultures of RSS916 (Figure 4G) than for GL (Supplementary Figure 7E) and WL grown cultures (Supplementary Figure 8E).

**Figure 4.**
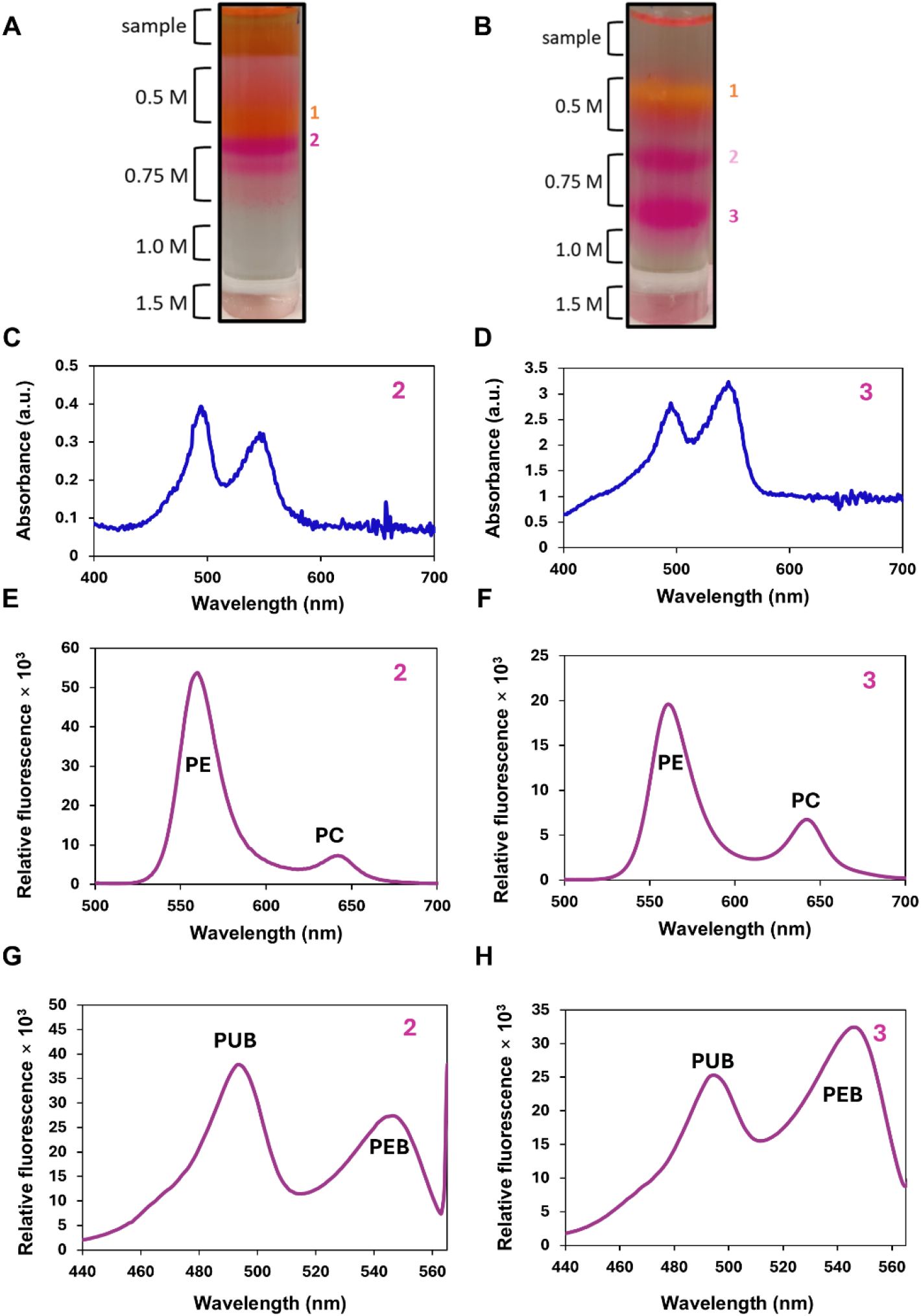
Isolated phycobilisomes from RS9916 and WH8109 grown in BL. Separation of the phycobilisomes on a discontinuous sucrose density gradient after 16 hr of ultracentrifugation of **(A)** RS9916 and **(B)** WH8109. UV-Vis spectra from 400 to 700 nm of **(C)** RS9916 fraction 2 on top of 0.75 M sucrose and **(D)** WH8109 fraction 3 beneath 0.75 M sucrose, exhibiting the PUB λ_max_ at 495 nm and the λ_max_ of 545 nm for PEB. **(E)** The emission spectrum of RS9916 fraction 2 and **(F)** WH8109 fraction 3, excited at 495 nm, recorded between 500-700 nm. A fluorescence excitation scan of **(G)** RS9916 fraction 2 and **(H)** WH8109 fraction 3, covering the range from 440-565 nm, with emission recorded at 570 nm. The fluorescence spectra for all samples were normalized to mg protein, as determined using the Bradford assay.

The PBS fractions obtained during sucrose gradient separation for WH8109 are presented for BL (Figure 4B), GL (Supplementary Fig. 9) and WL (Supplementary Figure 10). The sucrose gradients also exhibited similar separation patterns to one another (Fig. 4B, Supplementary Figure 9A and 10A). The yellowish pinkish fraction at 0.5 M, labelled as Fraction 1, was identified as Chl-*a*, due to its strong fluorescence at 680 nm after excitation at 440 nm (Supplementary Figures 9B and 10B). Fraction 2, at 0.75 M sucrose exhibited a PE associated fluorescence peak at 545 nm upon excitation at 495 nm, indicating that this fraction did not contain whole PBS, as PC was missing.

Fraction 3, located below 0.75 M sucrose, exhibited a UV-VIS spectrum with absorption peaks at the PUB λmax of 495 and the λmax of 545 nm for PEB (Figure 4D, Supplementary Figures 9C and 10C). The PUB peak was lower than the PEB peak under all three light conditions tested. Excitation of fraction 3 at 495 nm yielded the distinctive fluorescent emission peaks for PE, at 545 nm, and PC, at 630 nm (Figure 4F, Supplementary Figures 9D and 10D), suggesting that this fraction contained complete PBS. The fluorescence scan of fraction 3, recorded at 570 nm, showed that the relative PUB:PEB ratio for the whole PBS obtained from WH8109 remains low for all light conditions tested (Figure 4H, Supplementary Figures 9E and 10E). Fraction 3, obtained under both GL and BL conditions, was confirmed to represent intact phycobilisomes by SDS-PAGE (Supplementary Figure 11) and MS (data not shown).

### Sequencing of the WH8109 CA4-B island did not reveal any mutations

The potential CA4 capability of WH8109 would be regulated by the CA4-B island, located between gene loci Syncc8109_0453 to Syncc8109_457, a total of 5046 bp. Genes within the CA4-B island of WH8109 (NZ CP006882.1) were sequenced to check for any possible insertions, deletions, or inconsistencies that would result in the loss of its ability to conduct CA4, as reported by (Lovindeer et al., 2021). Three primer pairs were designed to cover the entire CA4-B island (Supplementary Table 1). The genes within this CA4-B island include the two transcriptional regulators, *fciA* and *fciB*, the lyases *mpeW*, *mpeU*, and *cpeZ*, as well as a gene of unknown function, *unk10* (Figure 1). An additional primer pair was made to amplify and sequence the *mpeQ* region (Supplementary Table 1), responsible for attaching the chromophore PEB to the specific cysteine residue of PE-II, thereby converting it to PUB.

The obtained sequences (Supplementary Data) were aligned with the WH8109_RC genome available on the NCBI under accession number NZ_CP006882.1. No insertions, deletions, or discrepancies were observed in the CA4-B gene clusters of WH8109 from our lab, indicating alignment consistency with the NCBI reference genome of WH8109.

## Discussion

Previous studies have reported conflicting results regarding the capacity of *Synechococcus* sp. WH8109 to undergo chromatic acclimation, prompting us to re-examine its response to changes in light colour. In the present study, we were unable to resolve the discrepancy between the findings of Humily et al. (Humily et al., 2014), who reported that WH8109 could not be induced to undergo CA under BL, and those of Lovindeer et al. (Lovindeer et al., 2021), who observed CA under the same conditions. We repeated the experimental conditions described by Lovindeer et al. (Lovindeer et al., 2021), including the use of SN medium, but were unable to induce CA4 in WH8109. To further investigate this discrepancy, we obtained a new WH8109 culture from the Roscoff Culture Collection, the same source used by Lovindeer et al. (Lovindeer et al., 2021), but again observed no CA4 activity. Thus, under the experimental conditions tested in this study, we found no evidence of CA4 activity in WH8109. *Synechococcus* sp. WH8109 maintained a fixed PUB:PEB ratio under BL, comparable to that of the GL specialist PT3a strains described by Dufour et al. (Dufour et al., 2025).

Sequencing of the CA4-B genomic island of WH8109 confirmed the presence of the characteristic regulatory genes *fciA* and *fciB*, as well as the phycobiliprotein lyase genes *mpeW* and *mpeQ*. Grébert et al. (Grébert et al., 2021) demonstrated that CA4- B involves the coordinated action of these two enzymes, which compete for the same chromophore-binding site (Cys83) of the α-subunit of PEII: under GL, MpeW attaches PEB, whereas under BL, MpeQ converts PEB to PUB, thereby enabling reversible adjustment of the PUB:PEB ratio. MpeW and MpeQ belong to the broader E/F family of phycobiliprotein lyases, with the corresponding CA4-A enzyme pair, MpeY/MpeZ, exhibiting the opposite division of activities, with MpeY functioning as a PEB lyase and MpeZ as a PEB lyase-isomerase (Fig. 5). The proposed evolutionary origin of CA4-B provides an interesting context for the presence of this system in WH8109. CA4-B is thought to have evolved from a PT3c-like BL specialist carrying a pre-existing MpeQ- like lyase-isomerase, with acquisition of *mpeW* and the associated regulatory components providing the additional capacity for GL-responsive PEB incorporation. Thus, the CA4-B island may represent an acquired regulatory and enzymatic module superimposed on a pre-existing BL-specialist state. Despite the presence of this complete CA4-B gene set (Figure 1), however, WH8109 failed to exhibit CA under the conditions tested here, suggesting that the presence of the genomic island alone is insufficient to ensure functional CA4. Whether the MpeW and MpeQ proteins of WH8109 indeed exhibit the activities proposed for CA4-B, and whether their regulation is functionally coupled to light quality, remains to be experimentally determined. However, based on their predicted quaternary structure (Figure 5), including the positioning of active site residues, we propse a bilin lyase function for MpeY and MpeW, while MpeZ and MpeQ are the corresponsing lyase-isomarase counterparts (Kumarapperuma et al., 2022).

**Figure 5.**
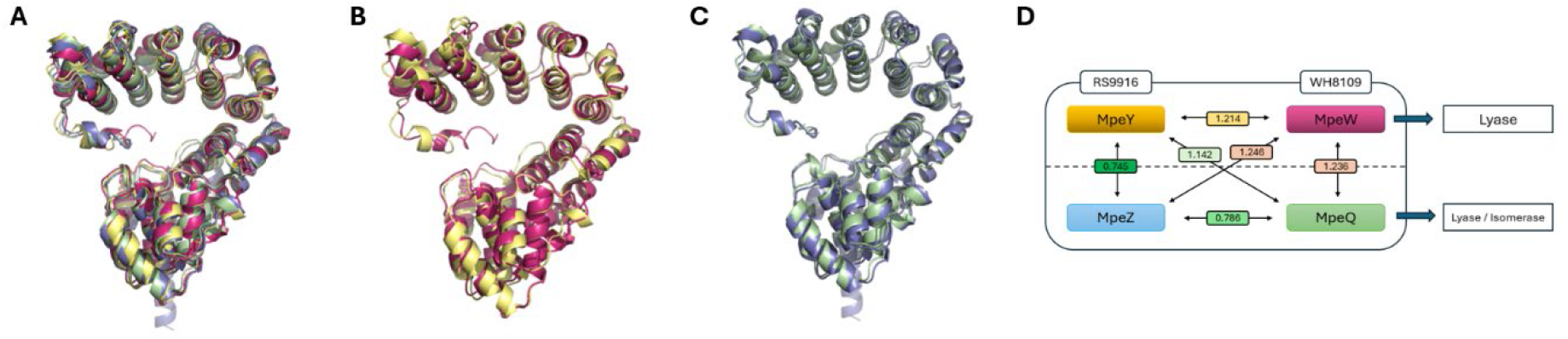
Structural comparison of Mpe proteins from *Synechococcus* strains RS9916 and WH8109. **(A)** Superposition of AlphaFold-predicted structures of MpeY (yellow) and MpeZ (blue) from RS9916, and MpeW (pink) and MpeQ (green) from WH8109. **(B)** Structural alignment of the Mpe lyases MpeY and MpeW. **(C)** Structural superposition of the Mpe lyase-isomerases MpeZ and MpeQ. **(D)** Pairwise root-mean-square deviation (RMSD) values for the individual protein structures. Lower RMSD values indicate greater structural similarity. Despite small differences between the proteins, the characteristic α-solenoid, question-mark-like architecture is retained, consistent with the structural organisation described by (Kumarapperuma et al., 2022).

This raises the possibility that the response of WH8109 to BL may depend on light intensity rather than spectral quality alone. The PT3a strain RS9907, for example, was unable to grow under low-intensity BL (15 µE), close to the 20 µE used here, but was able to grow at 75 µE (Dufour et al., 2025). Whether higher BL intensity can induce CA4-associated responses in WH8109 therefore remains an interesting question for future investigation, including possible induction of the lyase-isomerase gene *mpeQ*. Recent work further highlights the complexity of light adaptation in marine *Synechococcus*, which can involve changes beyond the PUB:PEB ratio, including PBS rod composition and the abundance and distribution of PBS, PSI and PSII complexes (Dufour et al., 2024; Dufour et al., 2025; Kis et al., 2026; Mattei et al., 2025). Dufour et al. for example, showed that the CA4-B strain RS9915 could alter its PE ratio under different BL and GL intensities, while the BL specialist RS9902 exhibited photophysiological characteristics that provided a competitive advantage under the conditions tested (Dufour et al., 2025). Such responses could provide alternative mechanisms for acclimation to changing light environments in strains that do not alter their PUB:PEB ratio.

For WH8109, photophysiological measurements under a range of light intensities could therefore help determine whether changes in antenna size, PBS composition or PSII efficiency accompany exposure to BL. Determining whether WH8109 alters its PE:PC ratio at higher BL intensity could indicate modulation of energy transfer along the PBS rods, as previously observed in other cyanobacteria (Stomp et al., 2004).

Overall, our results highlight that the presence of a CA4 genomic island does not necessarily correspond to detectable CA4 activity. WH8109 therefore provides an example of the distinction between the genomic capacity for CA and its phenotypic expression, and further work is needed to determine how light intensity and other photophysiological factors influence this relationship.

## Acknowledgements

We are thankful to Debbie Lindell (Technion, Haifa, IL) for the gift of *Synechococcus* sp. WH8109 and for training MF in cultivating it. We would like to thank Zuzana Storchova and Thorsten Stoeck (Faculty of Biology, RPTU), Kaiserslautern, Germany) for granting access of their flow cytometer instruments, Frederic Sommer and Michael Schroda for their help with mass spectrometry. This work was in part funded by the priority program SPP2330 of the German Research Foundation and the Landesforschungsinitiative BioComp to NFD, MF was funded by the ERASMUS+ program for his visit in Israel.

## Abbreviations

APC: allophycocyanin
APS: ammonium persulfate
BL: blue light
BSA: bovine serum albumin
CA: chromatic acclimation
Chl *a*: chlorophyll *a*
Em: emission
Exc: excitation
Exc495: ratio of excitation intensities at 495 and 545 nm
FI: fluorescence intensity
GL: green light; λmax wavelength of maximum absorption
µE: micro Einstein
PBS: phycobilisome(s)
PCB: phycocyanobilin
PC: phycocyanin
PEB: phycoerythrobilin
PE: phycoerythrin
PBP: phycobiliprotein(s)
PUB: phycourobilin
PUB: ratio of phycourobilin to phycoerythrobilin
WL: white light.

## Supplementary infromation

### Supplementary Figures

**Supplementary Figure 1.**
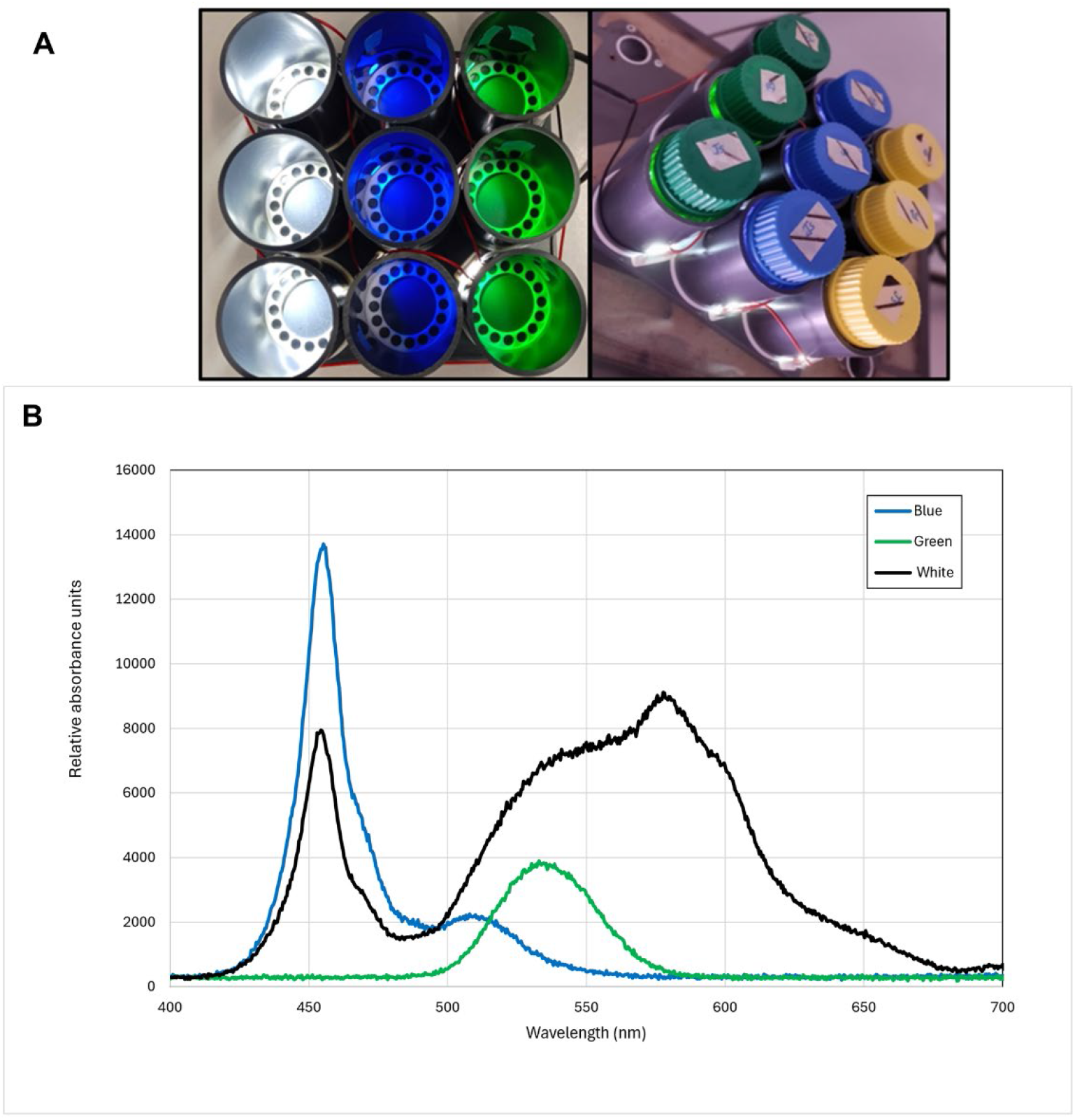
Apparatus used to culture WH8109 and RS9916, and the experimental light spectra used in this study. A) The culture set-up allowed individual lighting for each culture bottle set at 20 µE on a 14/10-hour light-dark cycle. The entire apparatus was placed on a shaker (120 rpm) in a culture room at 21 ± 2°C, Normalized emission spectra measured for the BL (dark blue, #119, 455 nm peak transmission), GL (primary green, #139, 525 nm peak transmission) and WL used to illuminate the cultures during acclimation experiments in the customised culture device used to grow *Synechococcus* strains WH8109 and RS9916.

**Supplementary Figure 2.**
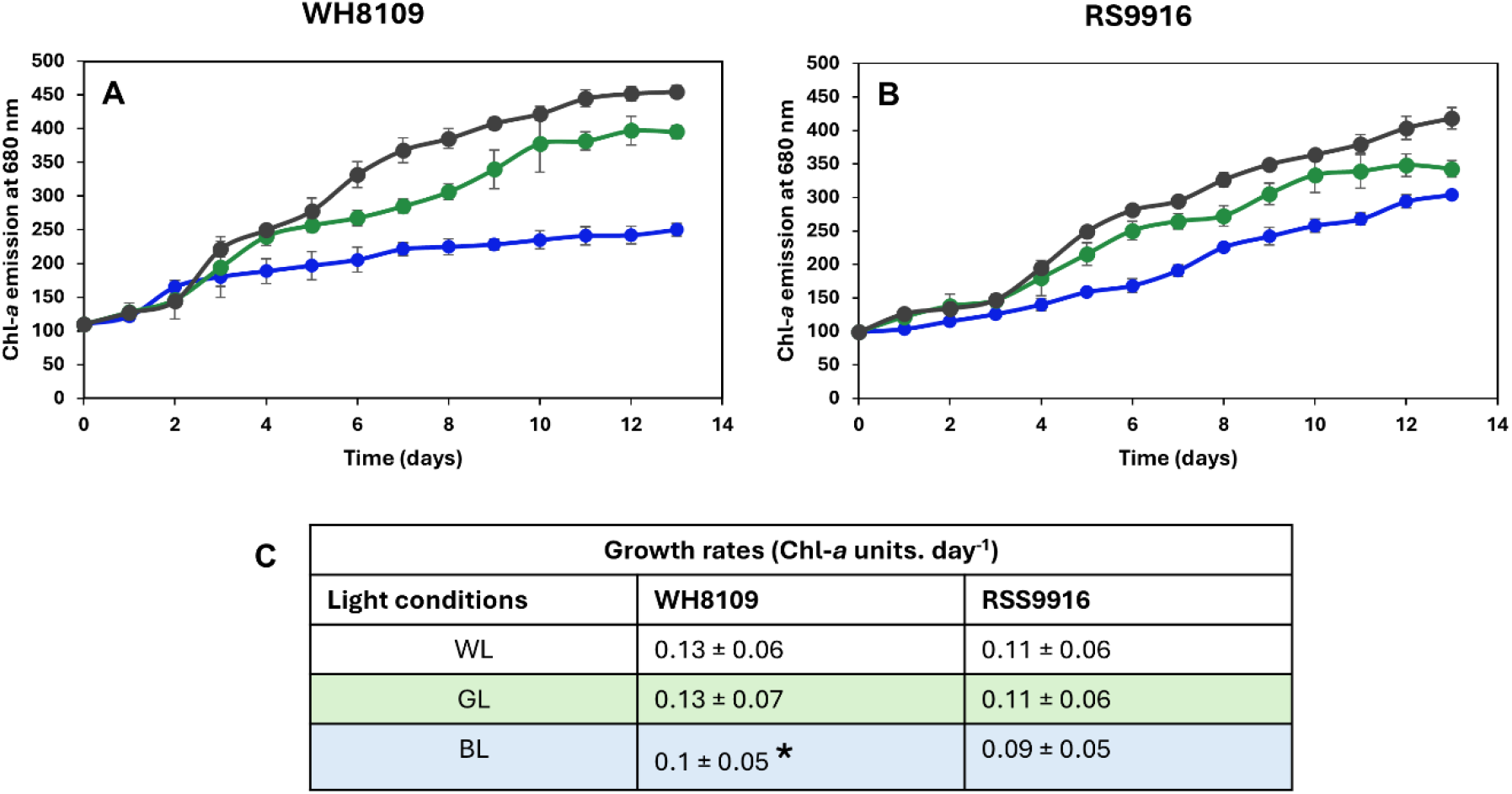
Growth rate of *Synechococcus* strains, WH8109 and RS9916 under blue, green and white light. Chlorophyll-*a* fluorescence emission recorded at 680 nm under excitation at 440 nm for (A) WH8109 and (B) RS9916. Blue lines represent the growth curve for cultures grown under BL, green lines represent GL grown cultures and black lines represent cultures grown under WL. (C) Table showing growth rates calculated from growth curves. The growth rates were calculated from the chl-a fluorescence units for the WH8109 grown under blue light (days 1 to 7), green light (days 2 and 10), and white light (days 2 and 11), for the RS9916 cultures cultivated under blue light (days 5 to 12), green light (days 3 to 10), and white light (days 3 to 12) (Supplementary datasets 1 & 2). All results are showing the mean ± SD of three independent biological replicates. Statistical significance was calculated between the WL, BL and GL conditions using a two-tailed Student’ t-test. Asterisk indicates significant differences to the WL grown cultures (* p ≤ 0.05).

**Supplementary Figure 3.**
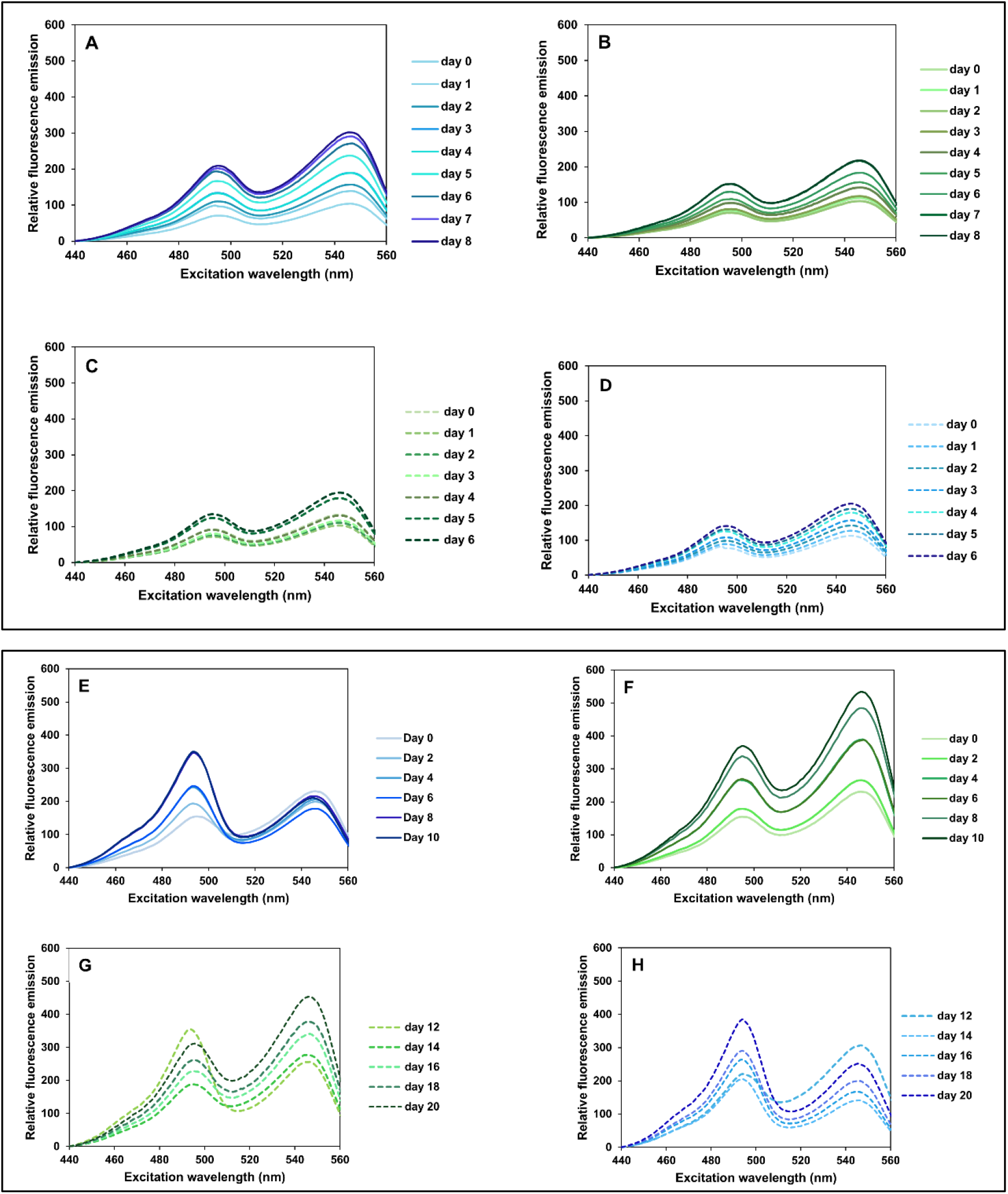
Emission spectra of whole cells of *Synechococcus* species cultivated in ASW medium and recorded through an excitation range from 495 nm and 545 nm. Plots of the 570 emission spectra recorded for WH8109 over an initial 8-day incubation period in BL (A) or GL (B). The culture material was subsequently diluted, and the cultures exposed to the opposing light conditions of GL(C) and BL (D), now plotted with dashed lines. Similarly, the emission spectra of for RS9916, the CA4 positive control strain, is presented for the initial 10-day incubation period in BL (E) or GL (F), with the ensuing emissions recorded after the cultures were swopped to the secondary light conditions GL (G) and BL (H), plotted with dashed lines.

**Supplementary Figure 4.**
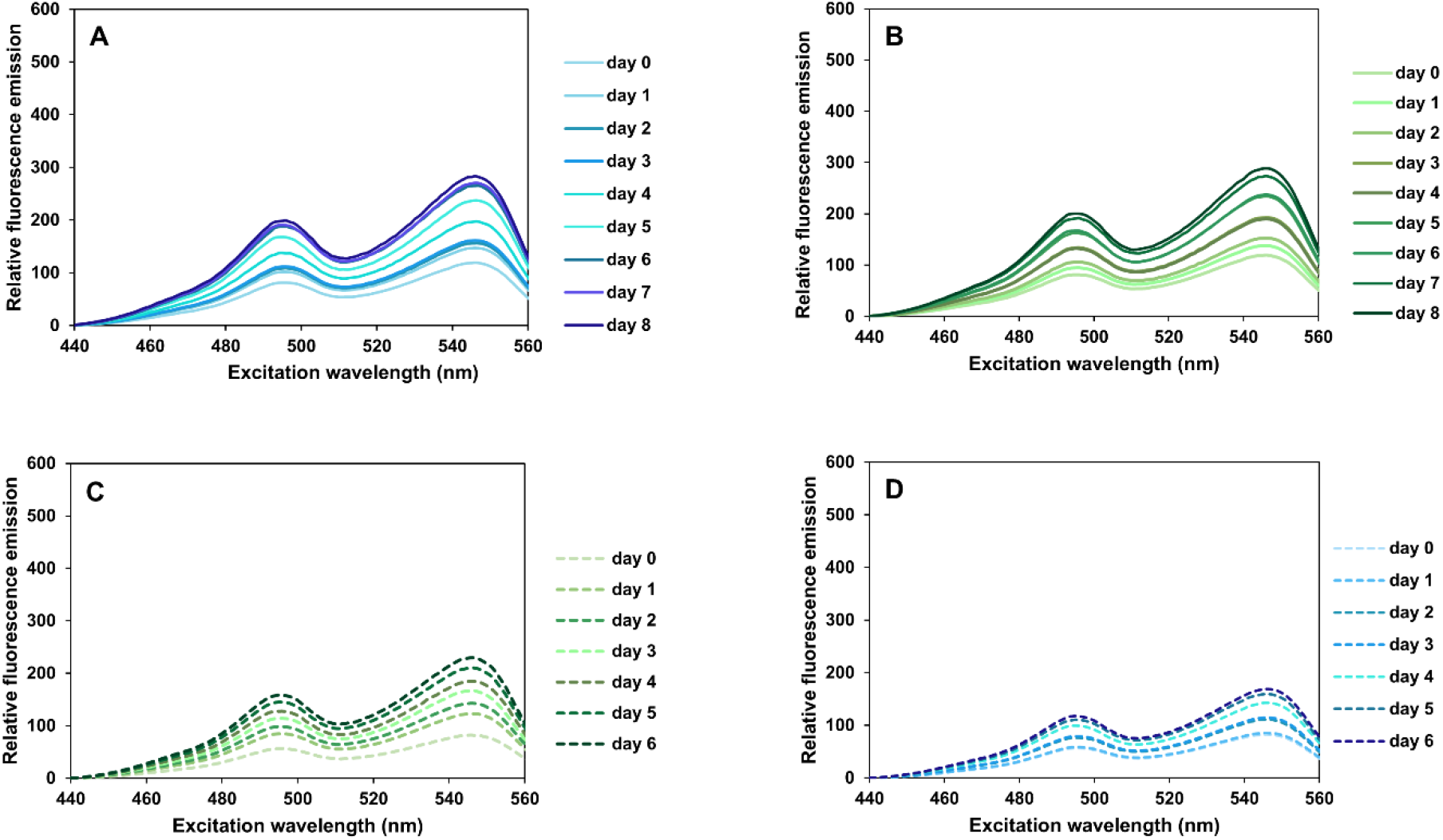
Emission spectra of whole cells of WH8109 cultivated in SN medium and recorded through an excitation range from 495 nm and 545 nm. To test whether the medium used influenced WH8109’s ability to conduct CA4, we repeated the experiment using SN medium (Lovindeer et al., 2021). Plots of the 570 emission spectra recorded for WH8109 over an initial 8-day incubation period in BL (A) or GL (B). The culture material was subsequently diluted, and the cultures exposed to the alternative light sources of GL(C) and BL (D), plotted with dashed lines.

**Supplementary Figure 5.**
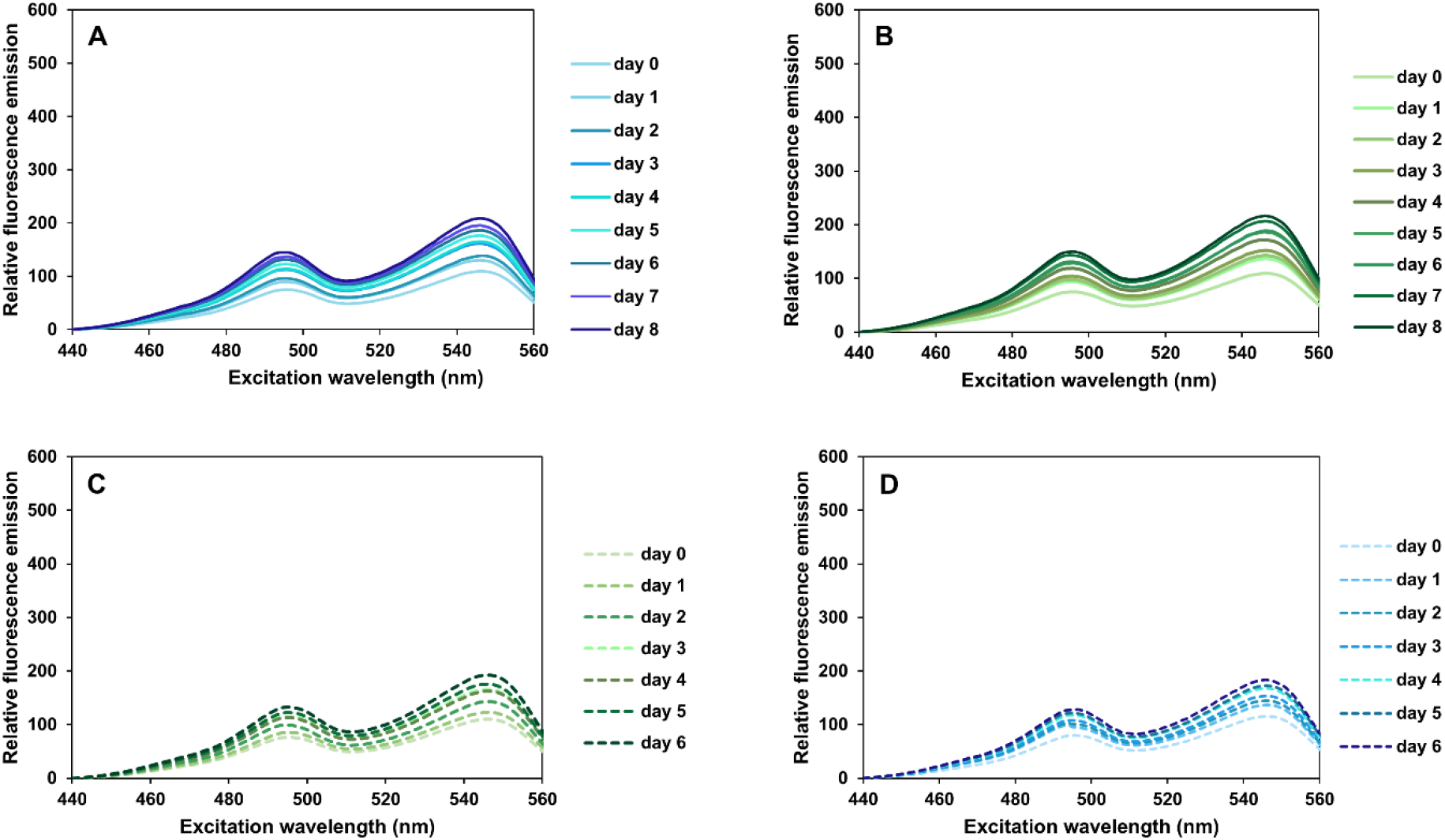
Emission spectra of whole cells of WH8109_RC cultivated in ASW medium and recorded through an excitation range from 495 nm and 545 nm. In order to test whether the original *Synechococcus* WH8109 strain conducted CA4, we repeated the culture experiments using a newly sourced culture from the Roscoff culture collection. Plots of the 570 emission spectra recorded for WH8109 over an initial 8-day incubation period in BL (A) or GL (B). The culture material was subsequently diluted, and the cultures exposed to the contrasting light conditions of GL(C) and BL (D), plotted with dashed lines.

**Supplementary Figure 6.**
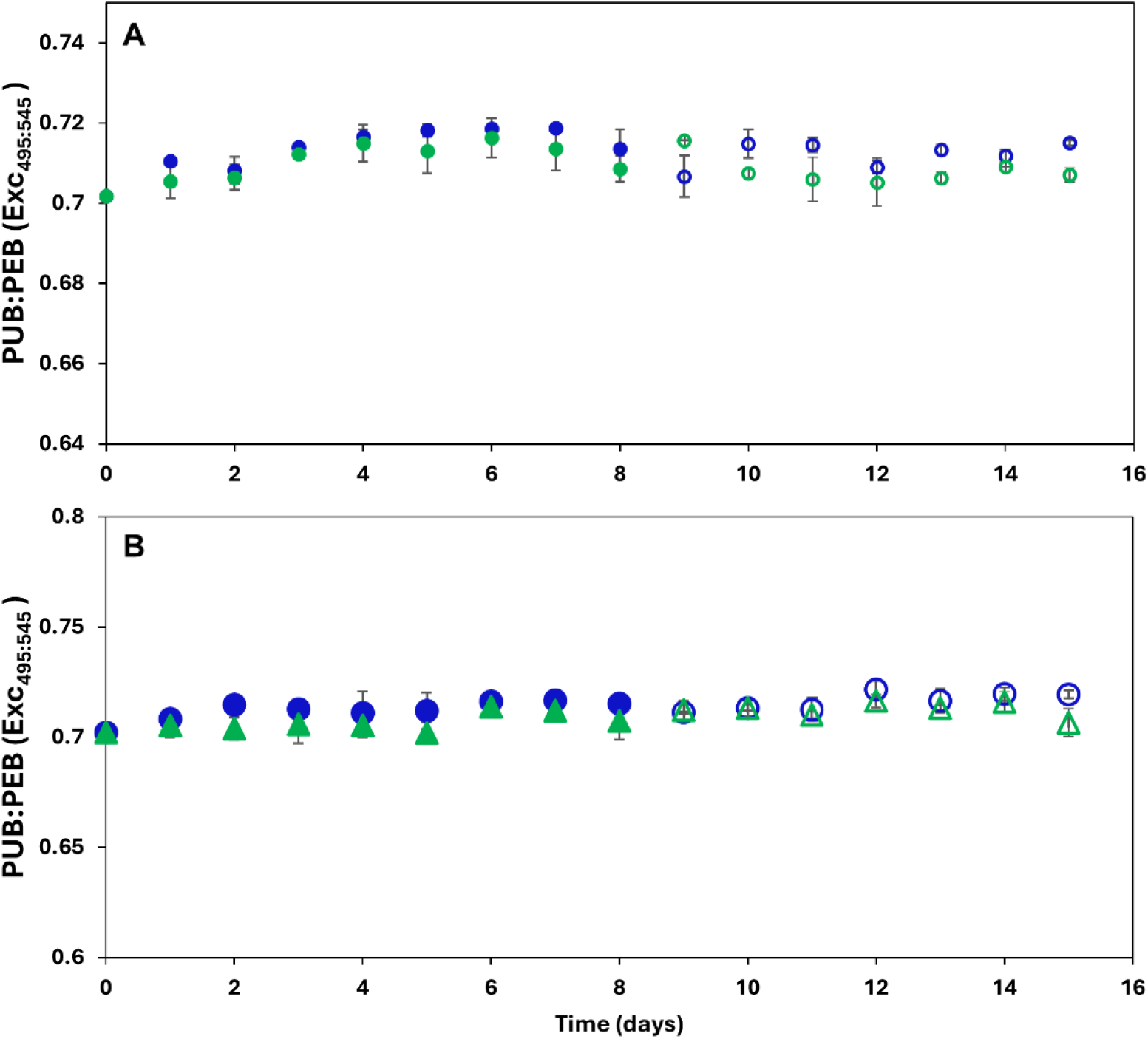
Comparative PUB:PEB (Exc_495:545_) *in vivo* fluorescence of *Synechococcus* WH8109 strains. Depicted are the average daily calculated PUB:PEB (Exc_495:545_) values during the light acclimation experiments for (A) WH8109 growing on SN medium and (B) WH8109_RC growing in ASW medium. Ratios obtained from the primary BL or GL growth phase (days 1-8) are plotted as blue or green dots respectively. Open circles represent the average PUB:PEB ratios in the cultures after the light was changed to the alternative light colour. Error bars represent 1 standard deviation (SD) of triplicate cultures and are often too small to be visible.

**Supplementary Figure 7:**
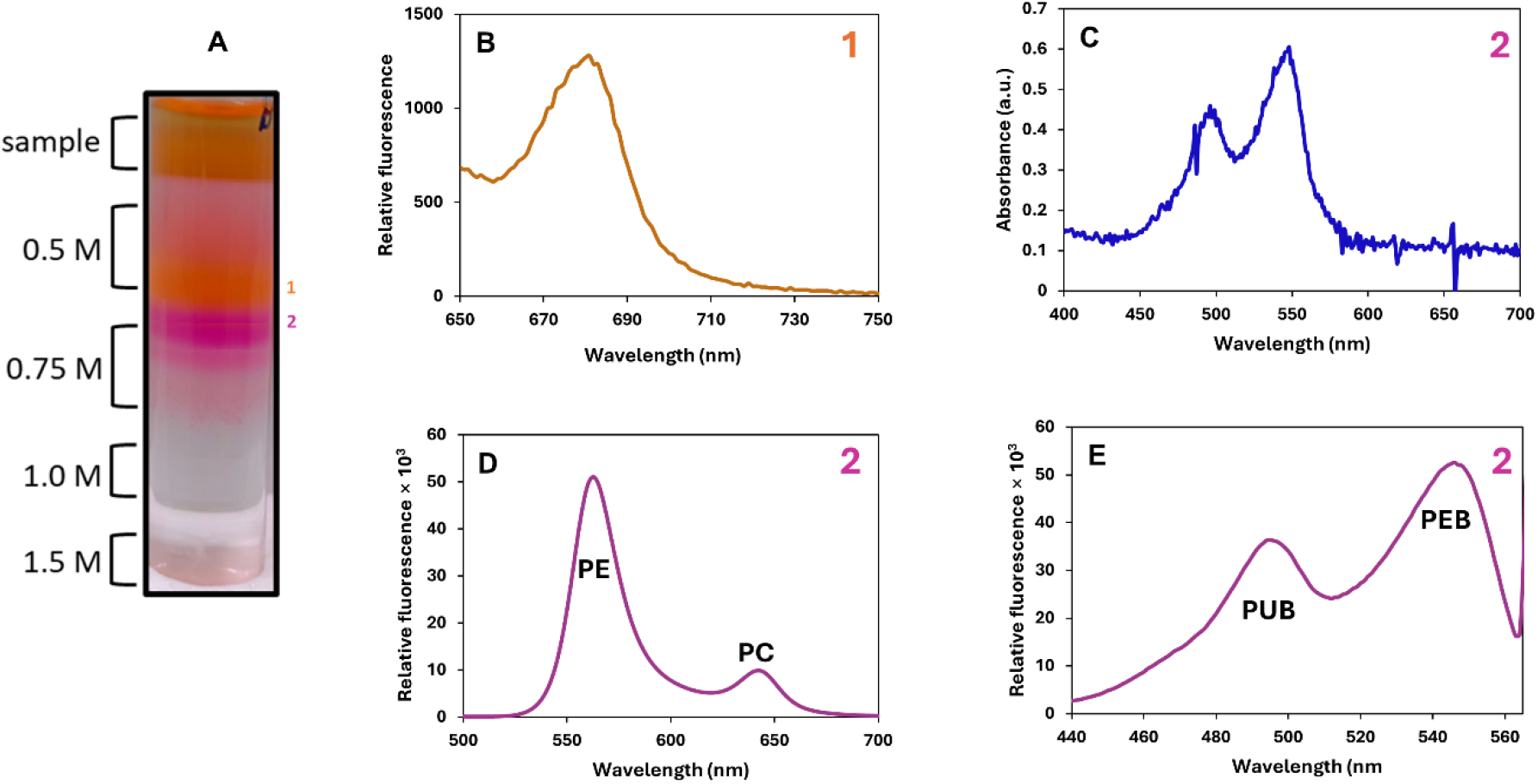
Isolated phycobilisomes from RS9916 cultures adapted to GL. (A) Separation of the phycobilisomes on a discontinuous sucrose density gradient after 16 hr of ultracentrifugation. (B) The emission spectra ranging from 650-750 nm of the first fraction at 0.5 M sucrose layer, with excitation at 440 nm. (C) UV-Vis spectra from 400 to 700 nm, of fraction 2 on top of 0.75 M sucrose, exhibiting the PUB λ_max_ at 495 and the λ_max_ of 545 nm for PEB. (D) The emission spectrum of fraction 2 excited at the 495 nm, recorded between 500-700 nm (E) A fluorescence excitation scan of fraction 2, covering the range from 440-565 nm, with emission recorded at 570 nm. The fluorescence spectra for all samples were normalized to mg protein, as determined using the Bradford assay.

**Supplementary Figure 8:**
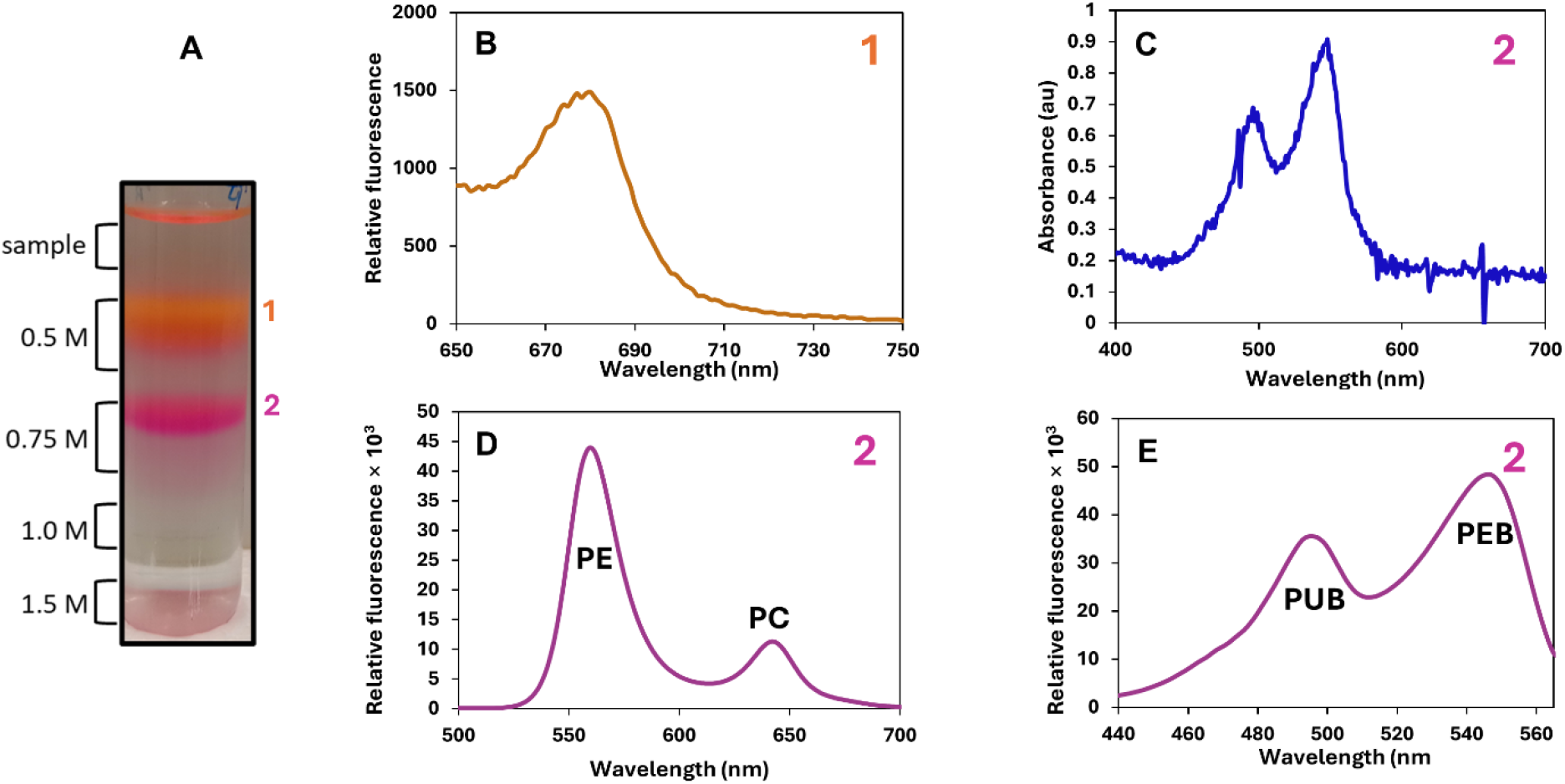
Isolated phycobilisomes from RS9916 grown under WL. (A) Separation of the phycobilisomes on a discontinuous sucrose density gradient after 16 hr of ultracentrifugation. (B) The emission spectra ranging from 650-750 nm of the first fraction at 0.5 M sucrose layer, with excitation at 440 nm. (C) UV-Vis spectra from 400 to 700 nm, of fraction 2 on top of 0.75 M sucrose, exhibiting the PUB l_max_ at 495 and the l_max_ of 545 nm for PEB. (D) The emission spectrum of fraction 2 excited at the 495 nm, recorded between 500-700 nm (E) A fluorescence excitation scan of fraction 2, covering the range from 440-565 nm, with emission recorded at 570 nm. The fluorescence spectra for all samples were normalized to mg protein, as determined using the Bradford assay.

**Supplementary Figure 9.**
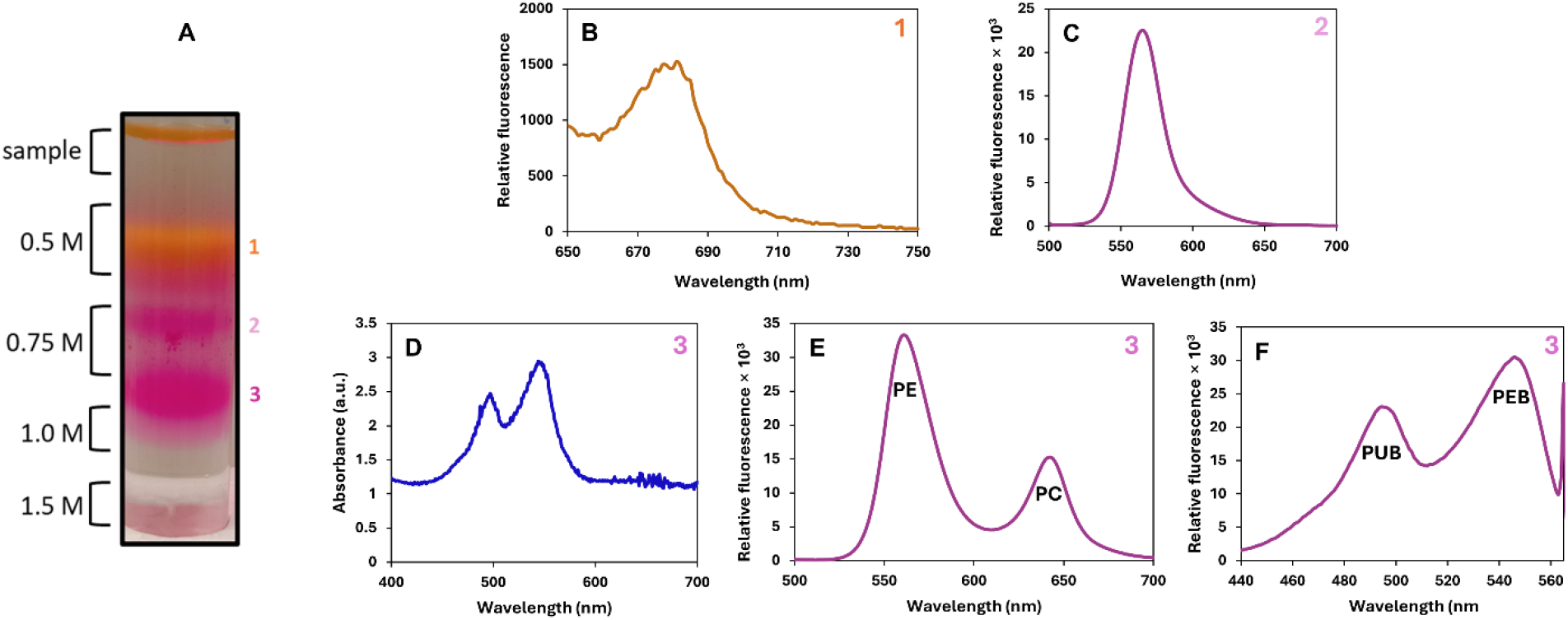
Isolated phycobilisomes from the WH8109 culture grown in GL. (A) Separation of the phycobilisome isolation on a discontinuous sucrose density gradient after 16 hr of ultracentrifugation. (B) The emission spectra ranging from 650-750 nm of fraction 1, with excitation at 440 nm. (C) The fluorescence emission spectrum recorded for fraction 2 from 500 to 700 nm, with excitation at 495 nm. (D) UV-Vis spectra of fraction 3, exhibiting the PUB λmax at 495 nm and the λmax of 545 nm for PEB. (E) The emission spectrum from 500-700 nm recorded by exciting fraction 3 at 495 nm. (F) A fluorescence excitation scan of fraction 3, covering the range 440-565 nm, with emission fixed at 570 nm. The fluorescence spectra for all samples were normalized to mg protein, as determined using the Bradford assay.

**Supplementary Figure 10.**
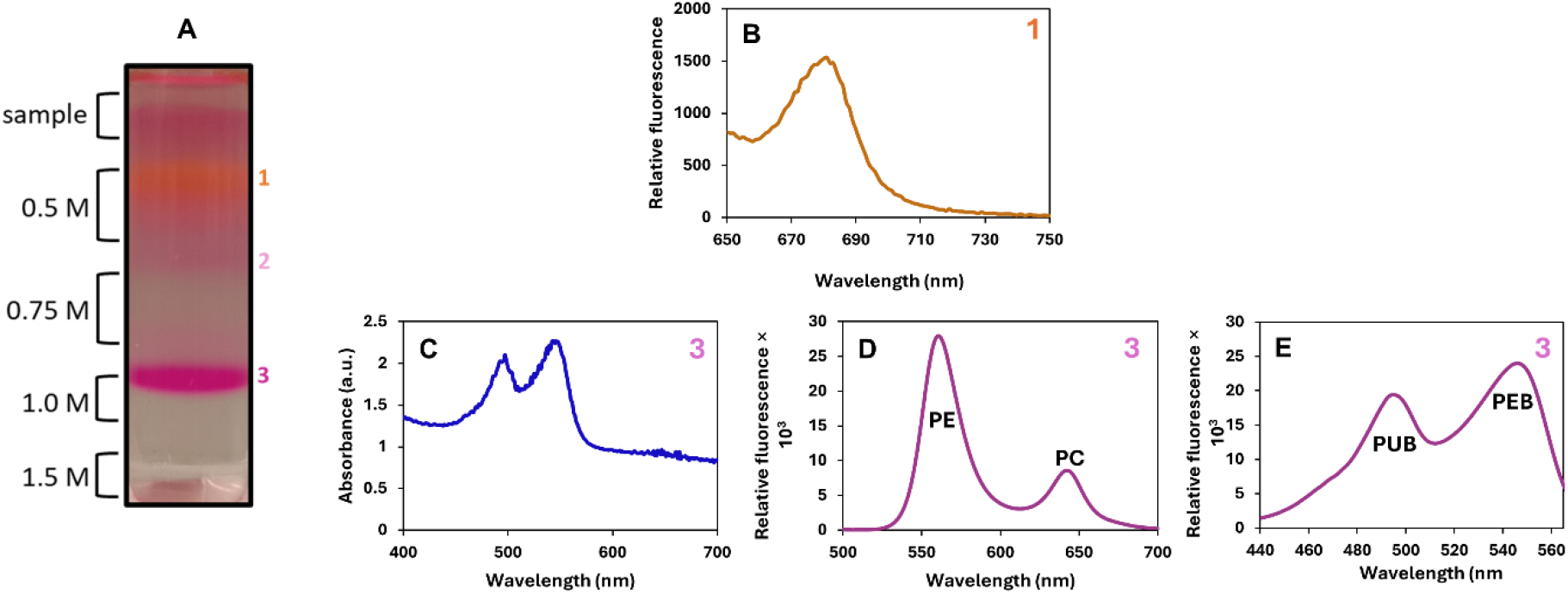
Isolated phycobilisomes from the WH8109 culture grown in WL. (A) Separation of the phycobilisome isolation on a discontinuous sucrose density gradient after 16 hr of ultracentrifugation. (B) The emission spectra ranging from 650-750 nm of fraction 1, with excitation at 440 nm. (C) UV-Vis spectra of fraction 3, exhibiting the PUB λmax at 495 nm and the λmax of 545 nm for PEB. (D) The emission spectrum from 500-700 nm recorded by exciting fraction 3 at 495 nm. (E) A fluorescence excitation scan of fraction 3, covering the range 440-565 nm, with emission fixed at 570 nm. The fluorescence spectra for all samples were normalized to mg protein, as determined using the Bradford assay.

**Supplementary Figure 11.**
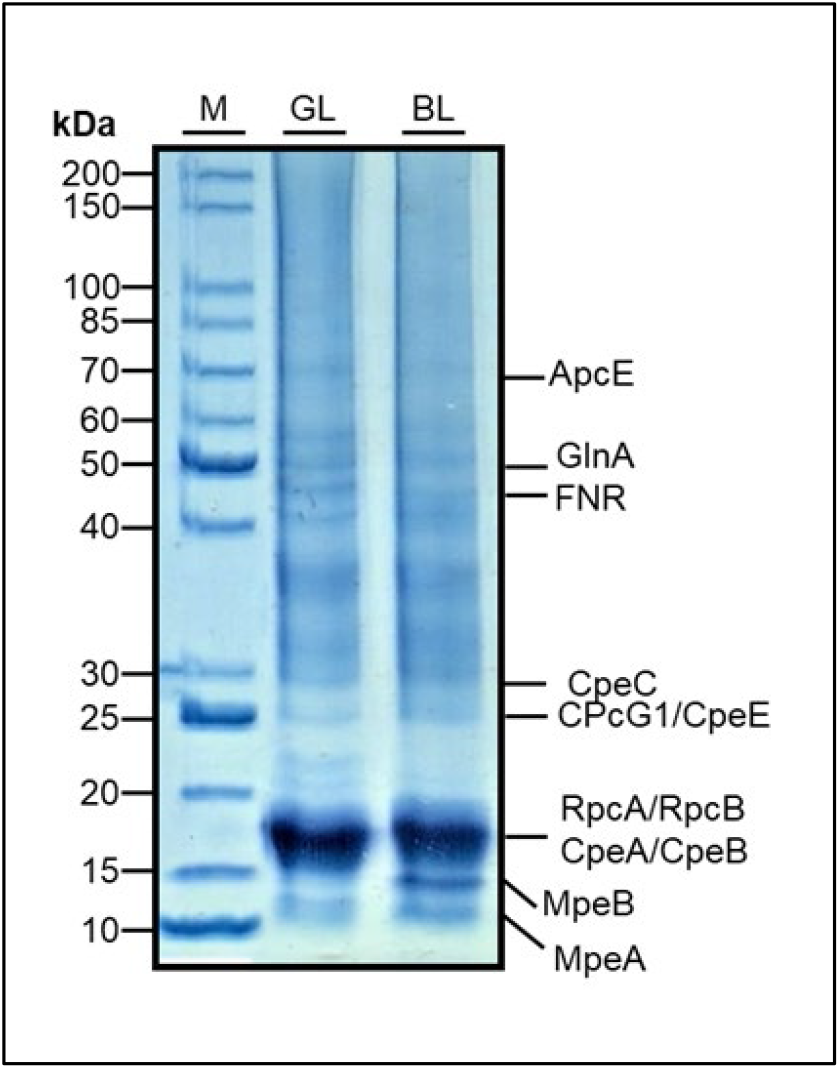
SDS-PAGE analysis of the intact phycobilisomes in fraction 3 from WH8109 grown under GL or BL. The proteins in the concentrated intact PBS fractions were separated by SDS-PAGE gel that was stained with Coomassie Blue for visualization. M = Unstained broad range protein marker (NEB P7717), with protein sizes indicated in kDa, GL = intact PBS from fraction 3 of WH8109 grown in GL, BL = intact PBS from fraction 3 of WH8109 grown in BL. Protein identification was confirmed by Mass Spectrometry analysis.

**Supplementary Table 1.**
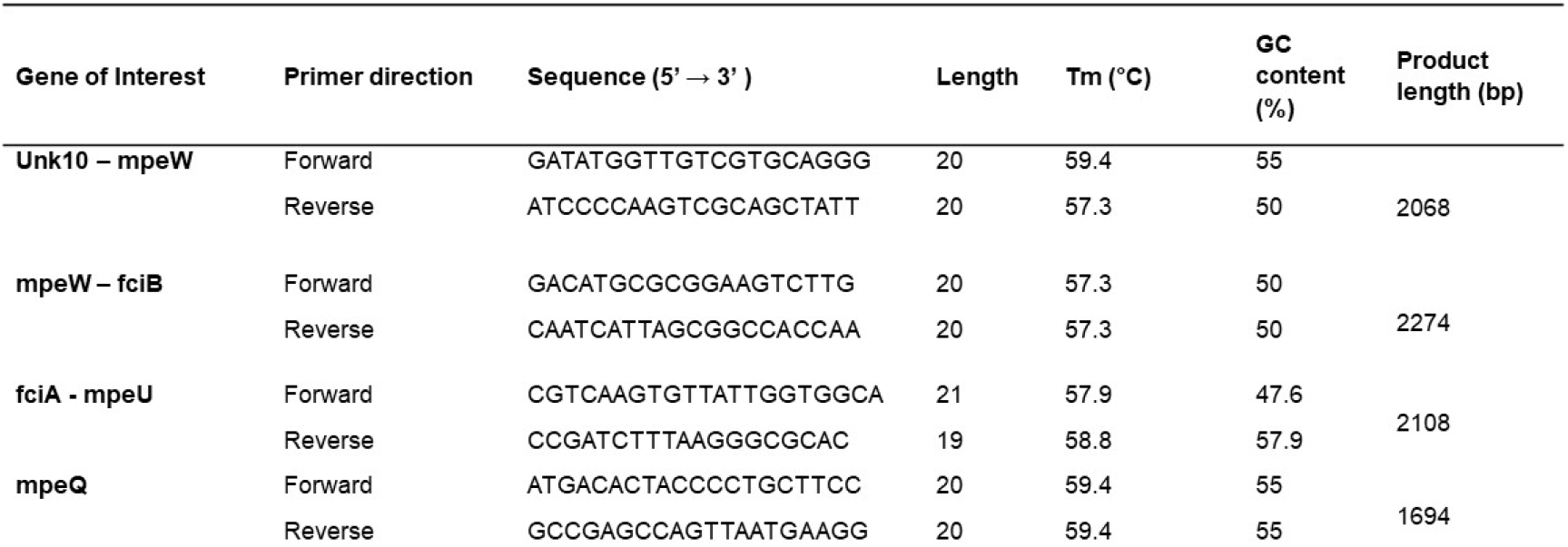
Oligonucleotide sequences used to amplify sections of the CA4-B island of WH8109.

### Supplementary Data

**Sequences of the genes associated with CA4 in the genome of WH8109 resequenced in this study** (original sequencing files are available upon request).

unk76.(Syncc810G_0453)

**345 bp**

TTATCGAGCTACAGTTTCTACACCTTTGTAGTAGCCTCCAGCGCGGAAATCAAAACGTTCTACTTTGTCTTCTTTTGCGCATGTTGTCTGTGAATAGTGCAGCAGAGTATTAAACCCTTCGGGCACCATCTTGGGGCACAGCCTCCCAACATCATTGGCAAATGGGCAGAAGTACACCCCTGCCCATCGATTAAGAAAACGTCCTTTTATTTCATCAGAAAAGATAAATCCATGTTGATTAGCGAGCTTGAGGATCTCCTCGGGCTCAGCTTCTTTGAAAAGGGATCTGAACGATAGGTCTTCTTCTAGAACCTGAAGAAAGCGGATAAACTCAGCACGAGACAT

mpeW.(Syncc810G_0454)

**1188 bp**

GTGCCAAGTTCACAGCGATTTGATAATCTTTTTTCATGGATGGGTGAACAGCAAGCAATTGAGTTGCTTTCGCAAAACGTTAGTACTCTTGATAATCCTGGCATTAAATATATAGCTGCAACACGGTTGGGAGCTTGTTCTTCACGAGATTCTTTGGATGCATTAGTCGCAGCGGCTACCGGGGATAGAGATAATATTTTCGATTCCATTACGCGTCGTAAGTCCATTGAAGCGCTAGGGCGTCGTCGCGACCTTAGTACTCTGCCAACCATTTATGAAGCTATGCATAGTAGTGATGAGCAGACGATTGTGAATTCAGTAGATACGCTAATTCATTTTGGCATACCTCTTGATGCTCAGTTTAAATCATCTCTTTTGAAGATTATTCAAAATGGCTCCGATATACTCAAGCGTGTTGCCATTCAATGCTTCACTCGTTTGGAGATGCATGAGTGTAATGGTATTATTCATGAGCAAGAGAGTAATCCAAATATATTGGTTAATGGCGCATCAATCGCTTATTCAATAAGGGTTGAAGATGACAAGCATAAATTGCATATTCTTGAAAATCATCTCAACGATTTGAATGTCATACATAGACGTTCGTCCGTCATTGATATTGGTGACGCTGGAGACCCTTCTCTTTTGTCAAGTATTTCGAAAGCAGCGGTGTCAATGCCGTTGAGAGCTAAAAGTGCGTTCAAGATGACTTCTAAAATTAAAACTAAATCACAAAGAACATTAATCAATCAAATGTTGAAGGATGATTCTCGACATTTGTATTTTACGCAGAAATTTGATGCACCAAATGATTTAGAAAAAGTTTGCGATTTCCTTAAGCATAGAGATGAAGAGCACCAGTACTTTGGTGCTAAGACCGTATTGAGTTGGCCAGCTTCAACTGTAACTGACGCAATTAAGTCTATTTGGGACCATCATG GATCTGATTATGGTGTTCACTATCAAGTAAATTGCTTGGTGTCTCAACTCGGCTTAAAAGAGTTGTCTTACATAATAAAGGAATCATTATTGGAGCCGGCTCCGCAGTATGCAAAATCAAAAATAGCTGCGACTTGGGGATGTTTAACTTTAGGTTTAATGGAATGTCGTAGTGATATTGAAAATCTACTTTTAAAATCCACTTGGGAGCCCCTACGATGGACATGCGCGGAAGTCTTGCGTGAACTA

fciB.(Syncc810G_0455)

**G66 bp**

CTATATTGGCAATCTATATCTTCTCGACTTTTCGATAGTCTGACTTGGGAGTTCTCCGAATGCAGTGTAATAGCGCCGTGCATATCTCCCTTGATGCTTAAACCCATATTTAATTGCAATATCTCGAATTGAATATAGATTTAGTTTTCGACGAGCGTCTTGGTTAAGCATAATTCTTCTGGATTCTTCAAGTCTAATTTGTGTCATCAATTCAATGATTCCCATTCCAAAGTACTCCTGACATATGCGATATAATGATGCTTGGCTTGTATTTATATATTTGCATACTTCAAGCAATGATAGAGGGGATTCCATCGTCTTCTCACTATGTGCAAGATCAATGATTTCTCTTAATTGATCGATATGTTTTGGATCTTGACTATTTTGCTCAGTCATTGGCTCTTCTAGACATGCGATTGCAAGATCAAAATATTTTTCTGGGTTCTGAATTCCCGCAGATACTTCTCTGGCAGCTAATCTCTTAAGTTGGTTTAGAGCTGTATTATTGCAATAAATCCCCTGATCACTCGTGAGTCGTTCAAGTCCATCATATGCTTTGCAATTTTCGAGTAATTCAAATAGGAGCTGCTTTTCTAGCATGCAACCAGTACTAGAACATTCGTTGTTATATTTTCCCCATGCGTTTCCTGGTACTGAATTAATCGTGCCAAATCCAGCCAGGCTATTTTTTGTCATTTTAAATCCATCACAACAGCCAAAGACTCCTTGAGTATTAGCTTTTGGCGTAATCCAATTGAAATCAATAGTTATACCATTCCCCCACCCCTCGTACAATAATGTTTTGTTGGCAGATATTTCTGCAAGTGCGACTTTATTAGTATTTGCTGCTTGCATAGTATAGAAACCACTGCCACCTTCCAGTTGCGTTATTTTTGTAAGCTGACCGAGTTTTGCAAGCTCTTCAGAGAGCTCCAGTGGATCACGATATTCAATAGATGTTTTCAT

fciA.(Syncc810G_0456)

**1005 bp**

TTACTCTCCCTGCCATGCACTAAATGAATACAAATTTTTATTGGATCGATCTATTGTTTGGCTGGGTGATTCCCCAAAGTATGTGGAGTAGGAGGCTTCGAATTTTTTCCAGTTCGTAAAACCGTAGTAAAGTGCTGTTCTCTTTTTTGTAAAATCTTTGAGACCTGTTGAGGAATGTGGAGTTAAAAAAGCCTTGCGAGTCTGTTCTAGGCGTACTCTTCTTACAAGCTCAATCACATCCATATTAAAGTTTTTTCGGCATATAGTATTGAGAGACTCTGTATTTGAATTCAAGTATTCAGTAATTTCGTCAAGTGTTATTGGTGGCATGCCAGCCCTATCTTCGTGAATTAGTCTAACCATGTCTTCTATTAGATCCGT TGACTCGCATCGATGTGCGTCATAGGTTGATTCATTATTTTCGTTGTCTAGGAATATAATTGCTAGATCGTAAAATGAATTTGGTTTCGTTGATGGCTGGCCATTAAAATGTCCGCTAAATAATTTTTTGAGCTGAGCACATGCATGATCATGCGAGTCAATTCCAATGCATTCCTCAAGCCTCGCATAGGCATTAAATGCGTTCAATTTGTTGATTCTCTCTTGTAATTTATTCCATTTCAGGCTCATGCAGCAAAGCTCTTCGTTTGCTCCTACAATGTCCCAGATATTTCCTCCTGTTTTGTTTATCCGGTTGAAGCCAGCAATGCTATCTTTTGTCATTTGGTGGCCGCTAATGATTGTCTGTGCCTCCGTATCGTTTGTCGGTAATGTGTCTAGCCAGCAGAATGATACTGATTTTGTGTTGGCTTCTTCCTCGTAGAGCAGAGTTTGGTTCGATTTAAAAATTTCAAGATGTACATCTTGAATAGGTGAGCAAGTGGTCTTCGTTTCGAGAAGGCCGGTGCTTAGTTGAGTTACATTGCATCCGAAGCTGGATGTTGAGTGATATTCCTCGAAATGTGCTTTGCAAAATTCCTCCATCTCGAGCGTATCCGAAAACAC

mpeU (Syncc810G_0457)

**8G7 bp**

TTATAAATCTCGGCTTAGTATATTAATTGCTGCATTGAATACTGCTTGGACTGACTCATCTGATTCTTTCGATTTCGCTTTTTCTATGCTTTTGATGCTGTTAATTGCCTCGAGCTTCATGAGTGCCATTGCTGTATTTTTGCGTACCTGAGTGTCTGTGTCGATTAATTTTTTGATAAGCATTTGTTCAACATCTTCGGCATCACATGATTTTCCTATGAGAGTTACCGCTTCTGCTCTCACCTCTGCTGAGATGTCATTGAGCGCTTGAAAGACTCTGTTTTGTGCCCGCAGATCATCACTCTTCTGTATTTGATCTCCCAAGGCTGAAATTGCAGCAACTCGAACTTCTGCCACGTCTGAACTGGCAGCCTCGAGCAATGCTTCCGGTGCTTTTGCGCCGATGAAACTCAGAGCAAGGTTGATCAAACCAACCTGGAATGGTGTGCAATCTGGATTTTTGAGGATTCCAAGCAACGAATCCATTGCGTCGGGGCCAATGGTTGCCATCGCTCCAGCAGACGACCCGAGTACAACTGGATCACTATCTTCTAAAAAGGCTTCAAGCAGATAAGGCAACGCGTCCTTGTTGCCAATCAAATTCAGTGTTTTGGCTGATGCTCTTCTTACGATCACATTCTGGTGCTGGCGAAGTGCTTTACAAAGAATGGGTAGAGCTTCGTTACCTACCGCCCCAAGACTTTTTGCAAAGGTGAGTCGTAAAGCGCCTCTCGCATCCCCAAGTCCGGCGACCATACGTTTGAGTGACTCGGCGTCAGATCTTGGGATCTCACCTTCGCTTAATTTTGCACTGAGTTCATCAGCGAGTTGGAGAGCTTCTTCTTCCGTGAGTTGGCTTTCGCTGATTGATGAGATATCGGTGAAATCCTGGAGCAC

mpeQ.(Syncc810G_0462)

**11G7 bp**

TGACAACTGTTTAAGGACGCGTTGGCAGGTCCATTTCAGTGGTAACCAGAAGGCACTGGCTGAAAGCTCCTCCAGGAGTGGCTTTTGGTCCACAAGGCCCAAGCGGAGGCACCCCCATGCAGCAGCGATCCGGGACTTGGTGTATTGAGGGATTGTTTCGGCTAAGGCAAGCCTGATTAGATCGCTTCGTTCTTCCAAACCCTGAAGACTGACGACGGCGGTTAGGTAATAGTGCGTAACGTAATCACTCCATAATTTTTCTTTAATTTCGTTAATTAAAATCATTCTCTCTGCTTTAGGCATGGCCATCAAGCTTGAGGCACCACCGTATTGTCGTGCTTCGTCGCGATGTTGAAGGTTGTTTTCGATCTCTGTAGGTTCAATTTTACAAACCCACTCTTTCCTAAGTTTGAGTTGTTGAGGGTTATCTTGCAAGAGTTGTGTTATTAGCTCAGCATATTCTTCCGGTATCTGGCATGTCTTATCAGGATCGACCAACTGAAATGCACTTCTAGCCCGCAAAGACATCGACACAGGAGCTGTGACAAGAGCTTCCAGCCTTGAAACGTCACCTGCATCGCCCAAATCAATTACGGCAGAGCGCCTCCGTCCCGCGATTGGATCAACAAGTTGCGGGATGAGGACATCAAGGCCCTCTGGCTGTTGAGCAACTTTCGACATGTATGCCCTTGCCGCACCAGCCACCAAGGGATTGGAATCATTCCGGAGGGGGCTGATGCTATTGATGACAGAGGGGACCCCTAAACGACAAAATGCTTGTATGACAGCTCGCTTTTGAATATCCTCACCCTCAAGGGCTTCCAATAATTTTTCATGATCTGCTTCTGTTAACGGTGCTCCAATTTTTGTTATCGCTTCTACAGAATTAATTACCGCAGCTTCATCGCTACATTTTAAGGCTCGAAACAGAGAAGGTAGGGCCTTTGGACTTTTTCGTCTGCCAAGGGCGTCAATCGCCTTGCGTCGCGTGATTCGATTAAAAAGATGCTCTGGATCCAGCGCTGCGGCCTGAATCAGCACATCAAGTGATTCTTCGCTGTTACTCGCTCCAAGCCTTGTGGCTGCCATGTATTTATCAACGGGCCTCTCGAGTGAGTCTGGATCAGCCAAAATCACTGCCATGGCCTGCTCTTCTGTCAAGC CTTCAACCAAGTTGTCGAATCGCTCGGCCAT

